# An inhibitor targeting glycosome membrane biogenesis kills *Leishmania* parasites

**DOI:** 10.1101/2024.09.13.612636

**Authors:** Shih-En Chou, Vishal C. Kalel, Ralf Erdmann

## Abstract

Leishmaniasis is a life-threatening neglected tropical disease caused by over 20 species of *Leishmania* parasites. Visceral leishmaniasis, also known as kala-azar, is particularly lethal, with a 95% mortality rate if left untreated. Currently, no vaccine is available, and chemotherapy remains the primary treatment option. However, these drugs have drawbacks such as high toxicities, the emergence of resistant strains, and high costs. Therefore, there is a need to develop new and safe treatments. Glycosomes are essential organelles for the survival of *Leishmania* parasites. They are maintained by peroxin (PEX) proteins, which are responsible for glycosome biogenesis, including targeting proteins to glycosomes. Previous studies have shown that blocking the interaction between the import receptor PEX19 and the docking factor PEX3 kills *Trypanosoma brucei* by disrupting glycosome biogenesis. In this study, we screened an FDA-approved drug repurposing library using an AlphaScreen based assay and identified inhibitors of *Ld*PEX3-*Ld*PEX19 interaction *in vitro*. The inhibitor effectively kills *Leishmania* parasites, including the challenging amastigote forms contained within the infected mammalian host cells. This study validates the inhibition of glycosome biogenesis in *Leishmania* as a potential approach for developing new anti-leishmanial therapies.

## Introduction

Leishmaniasis is a vector-borne neglected tropical disease (NTD) caused by infections of more than 20 *Leishmania* species. The clinical manifestations range from self-limited skin lesions at the infection area to disseminated mucocutaneous lesions or infections in internal organs, such as the spleen, liver and bone marrow (1–3). According to the clinical symptoms, the diseases are classified into three forms: cutaneous (CL), mucocutaneous (MCL), and visceral leishmaniasis (VL). The CL form is most common with skin sores developed over time after infection. Even though the sores are usually painless and might heal on their own without treatment, they leave disfiguring scars on patients and cause severe disability or stigma (4, 5). MCL, a severe type of CL, results in the destruction of oral, nasal, pharyngeal, and laryngeal mucosa and can lead to airway obstruction and aspiration pneumonia (4). VL, also known as kala-azar, is the second most fatal parasitic disease after malaria and one of the most threatening NTDs (6). The symptoms of VL include persistent fever, weight loss, anemia, and enlargement of the spleen and liver. Moreover, a sequel of VL, post kala-azar dermal leishmaniasis (PKDL), was recognized and clinically separated as a different form from CL since 1922 (7). It is characterized by macular, maculopapular, and nodular lesions on the face, upper arms, and trunk, and it usually happens 6 months to years after VL has been cured. Approximately 8% of patients who recovered from VL develop PKDL, but some cases of PKDL were reported without a clinical history of VL (8). Despite low fatality, patients with PKDL are thought to act as a reservoir for parasites because parasites are incubated in their skin (9). To date, about 6 to 12 million people are suffering from leishmaniasis across 90 countries (10, 11) and the infections mainly affect those in poverty, compounded by malnutrition and poor housing conditions, in Africa, Asia, and Latin America (3). Specifically, 50,000 to 90,000 new cases of VL occur annually (12). The other two NTDs, human African trypanosomiasis (HAT) and Chagas disease, also threaten humans in the world. HAT, caused by infection of *Trypanosoma brucei,* which tsetse flies transmit, is under control this decade with continued efforts (13). Around 6 to 7 million people worldwide are affected by Chagas disease after being infected with *T. cruzi*, which *is* spread by *Triatominae* (14). These NTDs, especially leishmaniasis, are significant health burdens that enormously affect medicine, society, and the economy.

Until now, no vaccine has been successfully developed to prevent leishmaniasis (15–17). Thus, chemotherapies are still used on patients despite their various disadvantages, such as different toxicity and high treatment costs (18, 19). Amphotericin B (AmB), miltefosine, paromomycin, and pentavalent antimonials (Sb) are mainly used first-line anti-leishmanial drugs. AmB can bind with ergosterol, an abundant component in fungal and leishmanial membranes, causing transmembrane pores and cell death because of electrolyte leakage (20–24). Moreover, AmB can mediate the production of reactive oxidative species (ROS), which might induce cell death (25, 26). Not only toxic to fungi and parasites, AmB is also toxic to human cells due to its cholesterol-binding properties. For improvement of bioavailability and cytotoxicity reduction, different forms of AmB have been developed. Monomer AmB shows good bioavailability but causes nephrotoxicity, and oligomers tend to accumulate in the liver and spleen (27). Combined with them, AmB aggregate is considered as a better form, releasing monomers over time, which can avoid immediate toxicity to the kidney and meanwhile reduce the accumulation in the liver and spleen (28, 29). Additionally, liposomal AmB formulations also help to reduce toxicity (30) and represent the best treatment for CL and VL, although its efficacy differs from patient to patient (29).

Another first-line drug, miltefosine, was originally developed for solid tumors (31, 32) and later used as an oral drug for leishmania therapy. The mechanism of action in miltefosine treatment is not fully elucidated, but some studies reveal that it can disrupt intracellular calcium homeostasis and effectively kill parasites (33). However, miltefosine has some disadvantages, such as low bioavailability and side effects. Patients suffering from leishmaniasis usually are also affected by malnutrition and malabsorption, which can lead to slow absorption and low drug bioavailability (34). Furthermore, side effects such as vomiting, diarrhea, dizziness, and fever were reported (35). Several forms of eye discomfort, including acute scleritis, corneal infiltration, and uveitis, were likewise reported during treatment (36, 37). The other first-line drugs, paromomycin, and pentavalent antimonials, also have some drawbacks. Paromomycin is a safer drug compared to the others but has low efficacy on CL and triggers resistance (38, 39). Pentavalent antimonials (Sb) have been used for the past decades and have caused an emergence of Sb resistance, especially in India (40, 41). Moreover, different toxicities of Sb, such as cardiotoxicity, nephrotoxicity, and hepatotoxicity, were reported (42–44). In summary, the challenges for combating leishmaniasis include ineffective therapies due to the emergence of drug-resistant strains and server-side effects. Therefore, the priority for leishmaniasis is to develop an effective, safe, and affordable treatment.

*Leishmania* parasites are transmitted to humans by the bite of infected female sandflies (45, 46). Two stages are involved in the life cycle of *Leishmania* organisms: promastigotes and amastigotes. The former, the developmental stage, proliferates in the sandfly midgut, and the latter, the immobile stage, multiplies in the macrophages of the host and infects other cells (47). Promastigotes are transformed into amastigotes and establish themselves as specific compartments called *Leishmania* parasitophorous vacuoles (LPVs) after they are phagocytosed into macrophages or neutrophils (48–50). Glycosomes are specialized peroxisomes found in these parasites. They compartmentalize the first seven enzymes of the glycolytic pathways and play roles in many metabolic pathways, such as glycolysis, purine salvage, and β-oxidation of fatty acids (51, 52). A group of proteins, peroxins (PEXs), are responsible for glycosome biogenesis, including the recognition and transport of proteins into glycosomes. PEX5, PEX7, PEX13, and PEX14 are involved in the glycosome matrix protein import machinery. PEX5 or PEX7 recognize and bind to the proteins with type1 or type 2 peroxisome targeting signal (PTS1/PTS2), then interact with membrane protein PEX13 and PEX14 to import proteins into glycosome (53–55). The glycosomal membrane protein import system is controlled by PEX19, PEX3, and PEX16 (55, 56). PEX19 serves as a cytosolic receptor, recognizing and binding to peroxisomal membrane proteins (PMPs) with membrane peroxisome targeting signal (mPTS). The membrane protein PEX3 serves as the docking site for PEX19-PMP cargo, wherein the cytosolic protein PEX19 interacts with PEX3, and PMP is inserted into the glycosomal membrane (55, 56). PEX16 is involved in the trafficking of PMPs from the endoplasmic reticulum to the peroxisome (57, 58). Glycosomes have been considered as a drug target for trypanosomiasis and leishmaniasis treatment because they are involved in many metabolic pathways (59). Disturbing matrix protein import with PEX14 inhibitors has been shown to be effective in killing *T. brucei* (60). Moreover, blocking the interaction between PEX3 and PEX19 will affect PMP insertion and subsequently disrupt the import of matrix proteins, impairing glycosome biogenesis and leading to parasite death. Previously, inhibitors targeting the PEX3-PEX19 interaction in *Trypanosoma brucei* were found to kill the parasites (61, 62).

In this study, we established an AlphaScreen-based assay with recombinant *L. donovani* PEX3 (*Ld*PEX3) and *Ld*PEX19 proteins. Further, we screened a drug repurposing library to find probable inhibitors for the PEX3-PEX19 interaction as anti-leishmanial drugs. Six candidate compounds were identified through screening, and five were confirmed to have leishmanicidal capacities. Among these candidates, Compound 1 was identified as a potential drug due to the killing efficacy on both *L. tarentolae* promastigotes and *L. amazonensis* amastigotes. Moreover, Compound 1 can disrupt glycosome biogenesis without affecting human peroxisome biogenesis. This study pharmacologically validates glycosome biogenesis as a drug target against leishmaniasis.

## Materials and methods

### Plasmid construction

*Ld*PEX3 and *Ld*PEX19 gene sequences were obtained from TriTrypDB (*LdBPK*_364210.1 and *LdBPK*_353310.1, respectively) and were amplified using the genomic DNA of *L. donovani* (ATCC 30030D, Lot no. 63292294) as a template. Primers RE7512 and RE7513 were used to clone *Ld*PEX3 without the predicted transmembrane domain (*Ld*PEX3Δ1-42) into a pGEX4T2 vector. *Ld*PEX19 was cloned into the pET21b and pET28a vectors using primer pairs RE7514-RE7515 and RE8162-RE8163, respectively. Primers RE8381 and RE8382 were used to insert a thrombin cleavage sit in the pET21b vector with *Ld*PEX19 (*Ld*PEX19-T-His6). The human gene sequences were obtained from GenBank. The *Hs*PEX3 gene (8504) without transmembrane domain (*Hs*PEX3Δ1-33) was amplified using primers RE8598 and RE8599 and cloned into pGEX4T2 vector using FastCloning (63). The *Hs*PEX19 gene (5824) was amplified by primers RE8653 and RE8654 and cloned into pET21b with a thrombin cleavage site. Detailed information of cloning strategies is shown in **Table 1**.

**Table 1.**
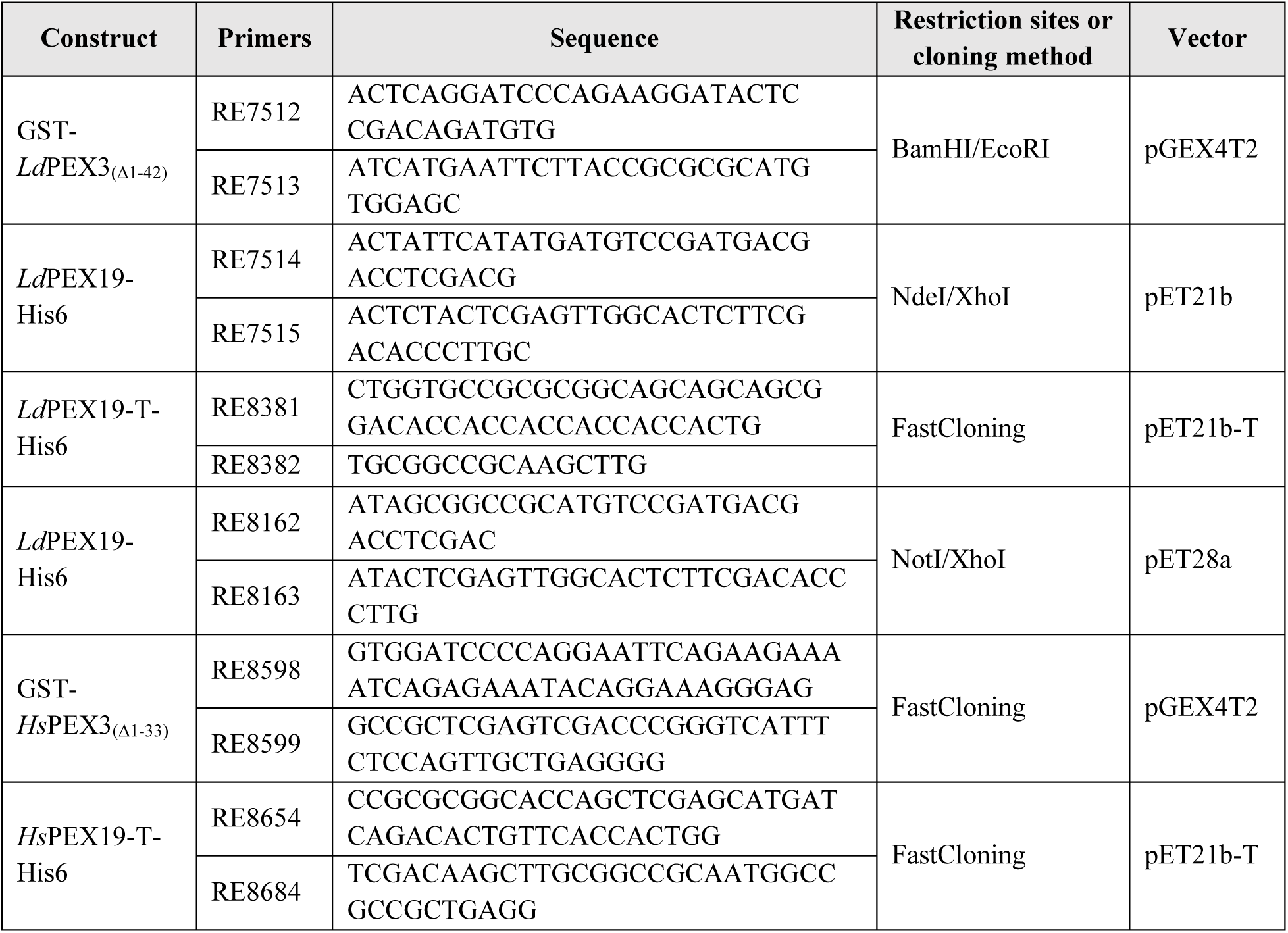

### Protein expression and purification

The plasmids mentioned in **Table 1** were transformed into *E. coli* strain BL21 (DE3) separately for protein expression, except for the *Ld*PEX19-His6 plasmids (in pET28a). Proteins with GST tags were expressed with 0.4 mM IPTG induction at 18°C for 16 h, and proteins with His6 tags were expressed with 1 mM IPTG induction at 30°C for 4 h.

All protein purification steps were performed at 4°C using cold phosphate-buffered saline (PBS, pH 7.4) as the buffer. Cells were suspended in the PBS buffer supplied with protease inhibitors (5 µg/mL Antipain, 2 µg/mL Aprotinin, 0.35 µg/mL Bestatin, 6 µg/mL Chymostatin, 2.5 µg/mL Leupeptin, 1 µg/mL Pepstatin A, 2.5 µg/mL DNAse, 1 mM PMSF, and 1 mM DTT) and mechanically lysed by EmulsiFlex (Avestin). The cell lysate was centrifuged in an SX4400 rotor (Centrifuge Allegra X-30R, Beckman Coulter) at 4,000 g for 10 min to separate cell debris and cell lysate. With high-speed centrifugation at 22,000 g in an SS34 rotor (Sorvall™ RC-6 Plus, Thermo Scientific) for 1 h, soluble and insoluble proteins were separated in the supernatant and pellet fractions, respectively. Subsequently, soluble proteins were purified by affinity chromatography on corresponding beads: Ni-NTA agarose beads (Macherey-Nagel) or glutathione agarose 4B beads (Macherey-Nagel). PBS supplemented with 200 mM imidazole or 10 mM reduced glutathione was used as the elution buffer for proteins with His6 or GST tags, respectively. The eluted proteins were dialyzed in PBS overnight using dialysis tubes (Spectra/Por, molecular weight cut-off 3.5 kDa). A Bradford assay (Coomassie Plus assay kit, Thermo Scientific) was performed to measure protein concentrations, and protein aliquots were stored at -80°C.

For co-expression, GST-*Ld*PEX3Δ1-42 and *Ld*PEX19-His6 (in pET28a) plasmids were co-transformed into BL21 cells and expressed with 0.4 mM IPTG at 18°C for 16 h. Subsequently, cell lysates with co-expressed GST-*Ld*PEX3Δ1-42 and *Ld*PEX19-His6 were purified as described above. Proteins were purified with glutathione agarose 4B beads using affinity chromatography. Recombinant GST-His6 complex encoded by pET42b plasmid was expressed and purified as described in (Li et al., 2021).

### Tag-free *Ld*PEX19 protein preparation

Recombinant *Ld*PEX19-T-His6 protein was used to generate tag-free *Ld*PEX19. 100 µg of recombinant *Ld*PEX19-T-His6 was incubated with 1 U of biotinylated thrombin enzyme (cat. No. 69672-3, Novagene) at 4°C overnight with gentle shaking. Then, Ni-NTA agarose beads were added to the reaction, binding to the cleaved His6 tag and any remaining uncleaved portion of *Ld*PEX19. The tag-free *Ld*PEX19 proteins were collected from the flow-through after loading beads into a Pierce™ Centrifuge Column (cat. No. 89896, Thermo Scientific). Tag-free *Ld*PEX19 was separated by SDS-PAGE and analyzed by immunoblotting with anti-His antibody.

### Establishment of an AlphaScreen based assay with recombinant *Ld*PEX3 and *Ld*PEX19

The Alpha (Amplified luminescent proximity homogeneous assay) approach is a beads-based technology to study interactions between molecules in a microplate. The assay relies on the binding between two different proteins or molecules to specific beads, and a luminescent signal is produced by an energy transfer from one bead to the other. In this study, AlphaScreen Nickel-chelate acceptor beads (cat. no. 6760619C, PerkinElmer) and AlphaScreen Glutathione donor beads (cat. no. 6765300, PerkinElmer) were used according to the tag in the recombinant proteins. The assay was performed in the 384-well plates (AlphaScreen-384 plates, PerkinElmer). Compared to the recommended Alpha beads concentration (20 µg/mL), we used a lower concentration (5 µg/mL) of Alpha beads as a final concentration in this study, which still gave robust and reproducible signals. For assay establishment, recombinant GST-*Ld*PEX3Δ1-42 and *Ld*PEX19-His6 proteins were cross-titrated. Proteins were tested at 8 concentrations with 3-fold serial titration from 0 - 300 nM. 5 µL of GST-*Ld*PEX3Δ1-42 were incubated with 5 µL of *Ld*PEX19-His6 and 5 µL of buffer, which is a substitute for a compound for 15 min. Then, 5 µL of acceptor beads were added to bind to *Ld*PEX19-His6 proteins in the dark for 15 min. Subsequently, 5 µL of donor beads were added to bind GST-*Ld*PEX3Δ1-42 proteins for another 40 min before reading the plate. Donor beads were excited with light at 680 nm, generating singlet oxygen molecules, which were transferred and reacted with acceptor beads. Subsequently, detectable light signals in the 520 to 620 nm range were produced and captured with a Cytation 5 plate reader (BioTek®) with the gain value set at 180. All incubation steps were performed at room temperature, and the above reactions were prepared in the PBS buffer supplemented with 0.5% BSA and 0.05% Tween 80.

Saturation and competition binding assay were applied to estimate the binding affinity between GST-*Ld*PEX3Δ1-42 and *Ld*PEX19-His6 proteins. In the saturation binding assay, three concentrations of GST-*Ld*PEX3Δ1-42 proteins (3 nM, 10 nM, and 30 nM) were separately incubated with different concentrations of *Ld*PEX19-His6. *Ld*PEX19-His6 was prepared with a 2-fold serial dilution starting from a maximum concentration of 300 nM, and a separate preparation of 200 nM of *Ld*PEX19-His6 was also made. In total, 11 concentrations of *Ld*PEX19-His6 were tested. For competition binding assay, recombinant GST-*Ld*PEX3Δ1-42 and *Ld*PEX19-His interacted at three combinations of concentrations, such as 10 nM-100 nM, 10 nM-200 nM, and 20 nM-200 nM. Further, 16 concentrations of tag-free *Ld*PEX19 with 2-fold serial dilution from a maximum concentration of 5000 nM were added to competitively bind GST-*Ld*PEX3Δ1-42 away from *Ld*PEX19-His6. Technical duplicates were performed in every experiment, and the means were used for analysis. Data for the saturation binding assay were obtained from three independent experiments. In the competition binding assay, results were collected from at least four independent experiments. Two types of curve fitting, one site-specific binding and one site-fit Ki in GraphPad Prism 10, were used for data analysis according to the needs.

### Repurposing library screening and hit selection in AlphaScreen based assay

The repurposing drug library (MCE-HY-L035P-30 PartA, MedChemExpress), including 3744 compounds, was tested in this study. Co-expressed and co-purified GST-*Ld*PEX3Δ1-42-*Ld*PEX19-His6 protein complex was used at final concentrations of 8 nM for primary and counter screens. 4 nM of recombinant GST-His6 protein complex was used as the final concentration in the counter screen to assess compounds interfering with the assay. Compounds from the library were screened at final concentrations of 10 µM in this study. 10 µL of GST-*Ld*PEX3Δ1-42-*Ld*PEX19-His6 complex was incubated with 5 µL of compounds for 30 min. Subsequently, acceptor and donor beads were added as previously described. Compounds were substituted by solvents (DMSO, ethanol, or water) as positive controls. Proteins incubated with high salt concentration (1 M NaCl) were used as negative controls. Buffer only with acceptor and donor beads was used to detect the background signal.

To inspect the screening quality, the Z-prime (Z’) factor was applied (64). The signals of positive (protein present) and negative (NaCl treated) controls were used for Z’ calculation. The Z’ factor above 0.5 indicated that the results from all the plates were good to distinguish between positive and negative controls. After confirmation with the Z’ factor, results from the plates were considered useable for interpretation. In each plate, alpha signals were transformed into robust Z-scores using median and median absolute deviation for calculation. A robust Z-score below -3 was set as the threshold for hit selection (65–67). The results fitting this criterion were thought to show significantly less signal than the positive control, and compounds were tested in the following experiments. Recombinant GST-His6 complexes were used as corresponding counter proteins for GST-*Ld*PEX3Δ1-42 - *Ld*PEX19-His6 complexes to exclude compounds that affect the protein tags or beads but not our interested proteins. 156 hit compounds were tested with recombinant GST-His6 and GST-*Ld*PEX3Δ1-42-*Ld*PEX19-His6 protein complex for the counter and the confirmation screens, respectively. The Alpha signals from the counter, and the confirmation screens were normalized to the positive control for comparison. Six compounds that showed specific inhibition of *Ld*PEX3-*Ld*PEX19 interaction were designated as true candidates and were further tested in the dose-dependent assay with different protein pairs using the AlphaScreen based assay.

In a dose-dependent assay, compounds were incubated with recombinant *Ld*PEX3-*Ld*PEX19 (8 nM), GST-His6 (4 nM), or pre-incubated *Hs*PEX3-*Hs*PEX19 proteins (10 nM and 30 nM, respectively) to estimate the inhibition effects. Thirteen concentrations of each compound from 0 to 100 µM with 2-fold serial dilution were tested. Data were obtained from at least three independent experiments with technical duplicates. Data analysis was performed in GraphPad Prism 10, with curves fitted using the dose-response-inhibition mode (inhibitor vs. response, three parameters).

### Enzyme-linked immunosorbent assay (ELISA)

The interaction between GST-*Ld*PEX3Δ1-42 and *Ld*PEX19-His6 proteins was tested using ELISA assay performed in 96 well microtiter plates (Immulon™ 2 HB 96-Well Microtiter EIA Plate, ImmunoChemistry Technologies) at room temperature. 1 µg of *Ld*PEX19-His6 was incubated in the well for 1 h, and unbound proteins were washed out three times with PBS. Then, 2-fold serially titrated GST-*Ld*PEX3Δ1-42, from 0 to 1000 nM, was added to each well and incubated for 1 h. In total, 16 concentrations of GST-*Ld*PEX3Δ1-42 were tested. Unbound GST-*Ld*PEX3Δ1-42 proteins were washed out with PBS. Subsequently, the bound GST-*Ld*PEX3Δ1-42 were detected by mouse monoclonal anti-GST antibody (Sigma-Aldrich, 1:1000), and the signal was amplified by anti-mouse horseradish peroxidase (Invitrogen, 1:1000). Substrate 3,3′,5,5′-tetramethylbenzidine (TMB, Thermo Fisher Scientific) was added following the antibody incubation. The reaction was stopped after 20 min with 0.5 M H2SO4, and the absorbance at 450 nm was monitored with a spectrophotometer. Data was obtained from three independent experiments with technical duplicates each and plotted with GraphPad Prism 10. The one site-specific binding model was used for curve fitting. The above reactions were prepared in the PBS buffer supplemented with 0.05% Tween20.

### Culture maintenance

*L. tarentolae* promastigotes (LEXSY host P10, Jena Bioscience) were used in this study. Parasites were maintained in Lexsy Broth BHI medium (Jena Biosciences) supplemented with 7.5 µg/mL hemin, 100 units/mL penicillin, and 100 µg/mL streptomycin at 28°C.

This study used the human cell lines T-REX293, HepG2 (Hepatocyte), GM05756 (skin fibroblast), HMC3 (microglia), and THP-1 (monocyte). T-Rex293 (Invitrogen) is a transformed primary human embryonal kidney cell line and was used to study the effects of compounds on human peroxisome biogenesis in the study. Other cell lines were used for cytotoxicity studies. All cells are maintained at 37°C with 5% CO2. All culture media were supplemented with penicillin (100,000 U/L), streptomycin (100 mg/L), and 10% fetal bovine serum (FBS), and different types of media were used depending on the cells. High-glucose Dulbecco’s Modified Eagle’s Medium (DMEM) supplemented with L-glutamine (4 mM) was used for T-Rex293, HepG2, and GM05756 cells. HMC3 cells were maintained in the Dulbecco’s Modified Eagle Medium/Nutrient Mixture F-12 (DMEM/F-12) supplemented with MEM Non-Essential Amino Acids Solution (MEM NEAA). THP-1 cells are cultured in Roswell Park Memorial Institute Medium (RPMI) supplemented with L-glutamine (2 mM). Before cytotoxicity testing, THP-1 cells were seeded in the 96-well plates with a density of 20,000 cells per well, and cell differentiation was induced by PMA (phorbol 12-myristate 13-acetate, 5 ng/mL) for 48 h. Subsequently, un-differentiated cells were washed out with HBSS, and differentiated THP-1 cells were tested for cytotoxicity.

### Assessment of anti-leishmanial activity

Candidate compounds were tested for the killing effect on *L. tarentolae* promastigotes using a resazurin-based cell viability assay. The assay has been widely used for the measurement of cell viability (68), based on the fluorescent signals from the reduction of non-fluorescent blue resazurin to a fluorescent dye (resorufin) by the mitochondrial respiratory chain in live cells (69). The assay was performed in 96-well plates. The tested compounds were prepared in 9 concentrations with 2-fold serial dilution starting from 200 µM. DMSO was used as a substitute for compounds as a negative control. 100 µL of parasite cells (5×10^4^ cells/mL) were added to the wells with 100 µL of prepared compounds or DMSO and then incubated at 28°C for 72 h. 25 µL of resazurin reagent (0.1 mg/mL in PBS (pH7.4)) was added to all wells following the 68-h incubation and further incubated for 4 h at 28°C. Following incubation, cell viability was estimated by measuring the fluorescence with a spectrophotometer (excitation 530 nm, emission 570 nm, and 585 nm). Wells with culture medium alone were used as a negative control for background signal, and untreated cells in the absence of compound were used as a positive control. Fluorescence values from 570 nm (as a reference wavelength) were subtracted from 585 nm, and the background signal was also subtracted. Following background subtraction, the positive control was set to 100% survival, and treated groups were displayed as percent survival values according to the positive control. Data was analyzed using a dose-response model (EC50 shift with X as concentration) in GraphPad Prism 10 to calculate EC50 (half maximal effective concentration).

### Compound cytotoxicity estimation in human cells

The cytotoxicity assay was performed in 96-well plates using a resazurin-based cell viability assay (68). Cells were seeded in wells and incubated at 37°C overnight. Cell density was dependent on cell lines. A density of 2000 cells per well was used for HepG2 and GM05756 cells, and a density of 5000 cells per well was used for HMC3 cells.

Tested compounds were prepared for 10 concentrations with 2-fold serial dilution ranging from 0 to 100 µM in culture medium. After removing the medium from the overnight cultured cells, fresh medium with compounds was added to each well and incubated for 72 h. After 68 h of incubation, resazurin assays were performed by incubating cells with resazurin for 4 h to determine cell viability. For GM05756 and HMC cells, 200 µM and 400 µM of Compound 1 were included for testing. Results were collected from at least three biological replicates with technical triplicates in each. Control (with 0 µM of compounds) was set up as 100% cell viability and used for normalization. Data was analyzed using curve fitting with a dose-response model (EC50 shift, X is concentration) in GraphPad Prism to calculate TC50 values.

### Leishmanicidal activity against *L. amazonensis* and cytotoxicity against host J774 cells

The *in vitro* leishmanicidal activity of compounds against *L. amazonensis* was performed by New York University (NYU) Langone’s Anti-Infectives Screening Facility in USA (on custom fee basis) to determine the EC50 of the compounds against intramacrophage amastigotes of *L. amazonensis* and the TC50 against J774 macrophage cells (70).

### Digitonin fractionation of *L. tarentolae* cells treated with Compound 1

*L. tarentolae* cells were treated with 12 µM Compound 1 in the density of 5×10^6^ cells/mL. As an untreated group (control), DMSO was added to cells with the same volume as Compound 1. The biochemical fractionation assay was performed as described in previous work (71). After 24-h treatment, cells were harvested and washed twice with PBS with 250 mM sucrose. The total protein was estimated using the Bradford assay, and digitonin was added according to the protein amount. The mislocalization of glycosomal matrix proteins was assessed with the immunoblot analysis using various antibodies such as GK (glycerol kinase), GAPDH (glyceraldehyde 3-phosphate dehydrogenase), and aldolase. Other antibodies, such as enolase (cytosolic marker), tubulin (cytoskeleton marker), and mtHSP70 (mitochondria Hsp70), were also used.

### Assessment of peroxisome biogenesis in T-Rex293 cells treated with Compound 1

T-Rex293 cells were used to test the effect of Compound 1 on peroxisome biogenesis. T-Rex293 cells with gene modification, such as PEX3 knockout (PEX3-KO) and PEX19-KO, generated previously in our lab (72), were used as a negative control to demonstrate that peroxisomal biogenesis was disrupted. Cells were seeded in 12-well plates (10^5^ cells) or on coverslips in 6-well plates (8×10^4^ cells) and incubated overnight. The next day, Compound 1 (12 µM) or DMSO (control) was diluted in the fresh medium and added to cells. Following 48-or 72-h incubation with compounds, cells in 12-well plates were washed twice to remove the medium, and cells were directly lysed by adding 1X Laemmli SDS sample buffer. Then, cell lysates were transferred to microtubes and denatured by heating at 95°C for 15 min. Samples were analyzed by immunoblotting using rabbit anti-ACAA1 (Thiolase) (1:2000, Sigma-Aldrich) and mouse anti-GAPDH (1:7500, Proteintech) antibodies. Cells in 6-well plates were used for immunofluorescence.

### Immunofluorescence Microscopy

*L. tarentolae c*ells were harvested, washed twice with PBS, and fixed with 4% PFA in PBS at room temperature for 15 min. Following fixation, cells were placed on a Poly-Lysine (10 µg/mL as the final concentration) coated slide for 1 h and permeabilized with 0.1% Triton X-100 in PBS for 10 min. PBS supplemented with 1% BSA and 0.25% Tween 20 were used as a blocking buffer and incubated for 1 h. Primary antibody rabbit anti-*Tb*Aldolase (1:500) was prepared in PBS and incubated with cells for 1 h. After washing five times with PBS, cells were treated with goat anti-rabbit secondary antibody (1:200, Alexa Fluor™ 594) for 30 min in the dark. After 3 times of wash, the stained samples were dried and mounted with Mowiol (Calbiochem, USA) containing DAPI (4′,6-diamidino-2-phenylindole). Samples were imaged with an Axio Imager M2 microscope (Zeiss, Germany) and analyzed using Zeiss Zen 3.2 software (blue edition).

T-Rex293 cells were washed twice using PBS and fixed with 4% PFA in PBS at room temperature for 20 min. Afterward, cells were permeabilized using 1% Triton X-100/PBS for 5 min and blocked with 1% BSA/PBS for 1 h. Following, cells were incubated with primary antibodies rabbit anti-PMP70 (1:500, Invitrogen) for 30 min and secondary antibody goat anti-rabbit secondary antibody (1:200, Alexa Fluor™ 594) for 10 min under light protection. The stained cells were mounted on glass slides with Mowiol supplemented with DAPI. Imaging was performed using an Axio Imager M2 microscope (Zeiss, Germany).

## Results

### Recombinant *Leishmania* PEX3 protein interacts with PEX19 protein in an AlphaScreen based assay

PEX3, a glycosomal membrane protein, and the cytosolic protein PEX19 play a key role in the import of glycosomal membrane proteins. Previously, we validated the PEX3-PEX19 interaction in *Leishmania* and found that expression of truncated PEX19 in *Leishmania* cells resulted in glycosomal morphology change (unpublished). Additionally, it has been shown that blocking the PEX3-PEX19 interaction in *Trypanosoma brucei* can kill the parasites (61, 62). Since these proteins are conserved within trypanosomatid parasites, the PEX3-PEX19 interaction is also expected to be a promising drug target for leishmaniasis due to its instrumental roles in glycosomal membrane protein import and glycosome biogenesis. In this study, we used an AlphaScreen based assay to assess the interaction between *L. donovani* PEX3 (*Ld*PEX3) and *Ld*PEX19 (the assay scheme shown in **Figure 1A**, **left panel**). Recombinant GST-*Ld*PEX3Δ1-42 and *Ld*PEX19-His6 **(Figure S1A and S1B)** were used for the assay establishment. Two proteins were cross-titrated at concentrations ranging from 0 to 300 nM to obtain the optimal protein concentrations, which produced a robust Alpha signal **(Figure S2A)**. Concentrations less than 11 nM of GST-*Ld*PEX3Δ1-42 did not give sufficient signals, and above 100 nM signal was saturated but did not show a hook effect till the highest concentration was tested, i.e., 300 nM. A hook effect indicates that the protein amount exceeds the binding capacities of beads, and proteins compete with each other to bind to the beads. Thus, the signal will drop when concentrations exceed the binding capacity of beads. Therefore, optimal concentrations can be chosen based on the hook point. Here, we observed a hook point at 11 nM of GST-*Ld*PEX3Δ1-42 and 100 nM of *Ld*PEX19-His6, so we chose the point with 11 nM of GST-*Ld*PEX3Δ1-42 and 33 nM of *Ld*PEX19-His6 before the signal reached the hook point. To simplify the handling, 10 nM of GST-*Ld*PEX3Δ1-42 and 30 nM of *Ld*PEX19-His6 were considered as optimal concentrations.

**Figure 1.**
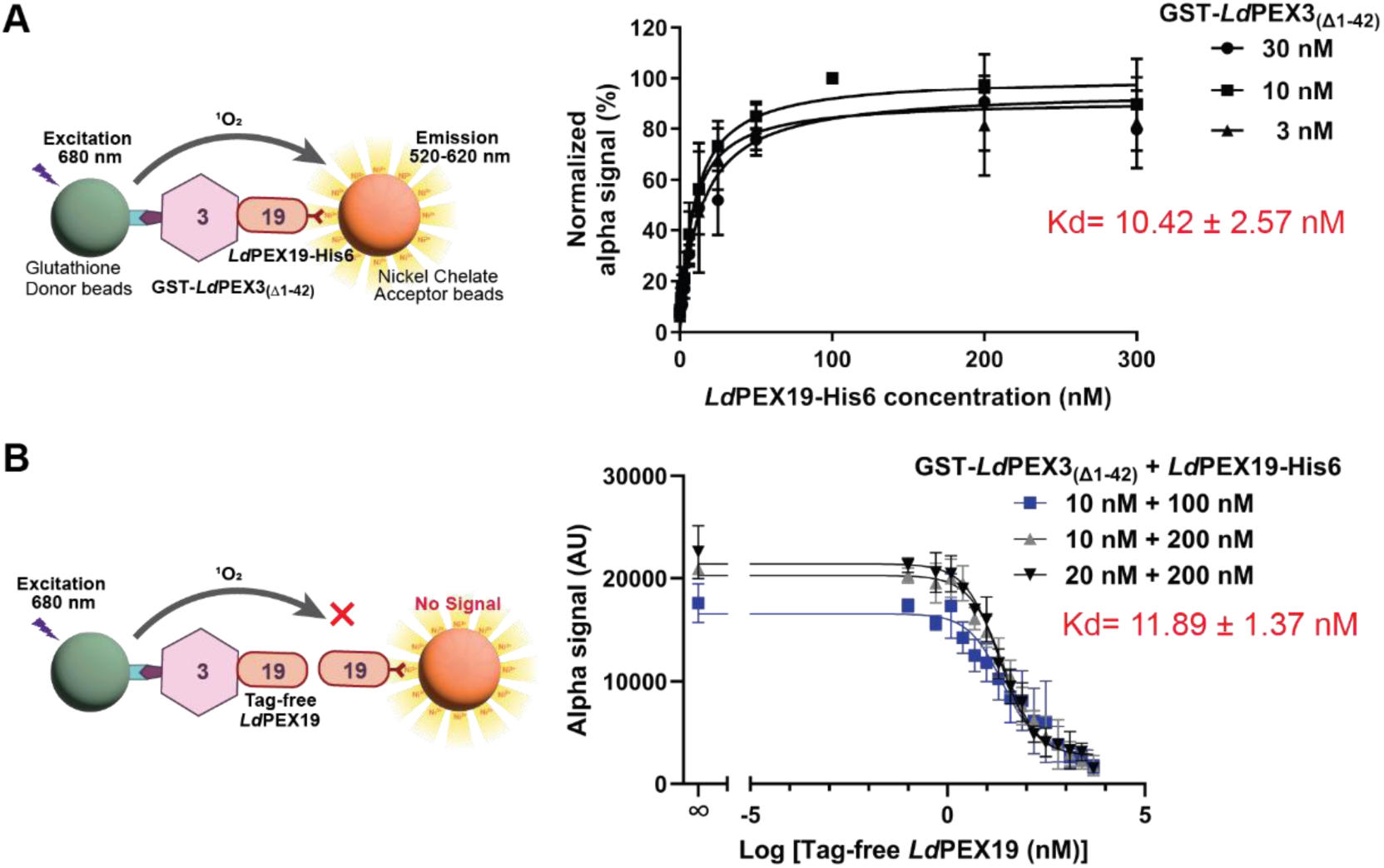
Establishment of AlphaScreen based protein-protein interaction (PPI) assay with *Leishmania* PEX3-PEX19 proteins. **A)** Schematic representation of the assay (left panel): The AlphaScreen based assay was established using recombinant GST-tagged *L. donovani* PEX3, which lacks the N-terminal predicted transmembrane domain (GST-*Ld*PEX3_(Δ1-42)_), and C-terminally His6-tagged full-length PEX19 (*Ld*PEX19-His6). Glutathione donor beads and nickel chelate acceptor beads decorated with the respective tagged fusion proteins were employed. When the proteins interact and thus bring the corresponding beads in proximity, singlet oxygen molecules are generated by the donor beads and transferred to the acceptor beads, producing a detectable light signal in the 520-620 nm range (left panel). The interaction between the proteins was analyzed using the saturation curve assay (Right panel). GST-*Ld*PEX3_(Δ1-42)_ was tested at three concentrations (3 nM, 10 nM and 30 nM) and titrated with 10 concentrations of *Ld*PEX19-His6 to assess interaction affinity. The results were collected from triplicate experiments under each condition and analyzed using one site-specific binding in GraphPad Prism 10. The dissociation constant (Kd) was estimated to be 10.42 ± 2.57 nM across the three conditions. **B)** Recombinant tag-free *Ld*PEX19 binds to GST-*Ld*PEX3_(Δ1-42)_, resulting in no detectable signal (left panel). In the competition assay, various concentrations of GST-*Ld*PEX3_(Δ1-42)_ and *Ld*PEX19-His6 proteins were allowed to bind, after which tag-free *Ld*PEX19 proteins were introduced to compete with *Ld*PEX19-His6. Reduced signals were observed when GST-*Ld*PEX3_(Δ1-42)_ bound to an excess of tag-free *Ld*PEX19, indicating that the *Ld*PEX3-19 interaction can be inhibited or competed. Data was collected from four or five replicates under each condition and analyzed using one site-fit Ki in GraphPad Prism 10. The Kd was calculated to be 11.89 ± 1.37 nM across the three conditions. Error bars represent the standard deviations.

According to the optimal concentrations determined above, we used three concentrations, 3 nM, 10 nM, and 30 nM of GST-*Ld*PEX3Δ1-42, in the saturation binding assay. The assay was performed to show the interaction between the proteins in the AlphaScreen-based assay. Different concentrations of *Ld*PEX19-His6 proteins were tested in this study, and the saturation assay results revealed the dissociation constant (Kd) to be 10.42 ± 2.57 nM **(Figure 1A**, **right panel)**. To assess the stability of *Ld*PEX3-*Ld*PEX19 interaction, a competition assay was performed using tag-free *Ld*PEX19 proteins. For this assay, recombinant *Ld*PEX19-T-His6 was purified by affinity chromatography on Ni-NTA agarose beads **(Figure S1C)**. Then, tag-free *Ld*PEX19 was obtained by removing the C-terminal His6-Tag from *Ld*PEX19-T-His6 by thrombin cleavage, and immunoblotting was applied to check the successful removal of the His6 tag **(Figure S1D)**. In the competition binding assay **(Figure 1B**, **left panel)**, signal reduction was observed when high concentrations of tag-free *Ld*PEX19 were added to compete with *Ld*PEX19-His6. The competition assay determined the Kd to be 11.89 ± 1.87 nM **(Figure 1B**, **right panel)**. Additionally, the ELISA assay was performed with recombinant *Ld*PEX3 and *Ld*PEX19, which similarly revealed a Kd of 10.63 ± 2.16 nM **(Figure S2B),** consistent with the results from the AlphaScreen based assay **(Figure 1A**, **B)**. Taken together, these findings demonstrate that *Ld*PEX3 interacts with *Ld*PEX19 and that this interaction can be competed/displaced with tag-free *Ld*PEX19 in the AlphaScreen based assay. This provides an optimal system to screen for inhibitors that block the *Ld*PEX3-*Ld*PEX19 interaction, serving as proof of concept.

### Screening and validation for inhibitors of *Ld*PEX3-*Ld*PEX19 interaction

After validating the interaction of *Ld*PEX3-*Ld*PEX19 in the AlphaScreen based assay, we utilized this assay to screen inhibitors of this interaction. To reduce handling steps during drug screening and for accuracy, recombinant GST-*Ld*PEX3Δ1-42 and *Ld*PEX19-His6 were co-expressed in *E.coli* and co-purified by affinity chromatography using GST beads **(Figure S1E)**. The isolated *Ld*PEX3-*Ld*PEX19 complex was also titrated to reveal the ideal concentration for the assay, which was 8 nM **(Figure S2C)**. Here, we utilized a repurposing drug library for the inhibitor screen, which contains FDA-approved and phase II-passed drugs. The screening was performed in 384-well plates, and 320 compounds from the repurposing drug library were tested in each plate. Each plate contained positive controls, which represented the protein binding without compound treatment, and negative control, which displayed the signal with a disrupted protein interaction when 1M NaCl was used. Beads only with buffer were used to show the background signal in the absence of either protein. The original Alpha signals of one screening plate were shown with controls in **Figure 2A**. To interpret the results from each plate, the Z-prime (Z’) factor was used to evaluate data quality. Z’ factor above 0.5 indicates the reliability of each screening plate in distinguishing negative and positive results. In total, 15 plates were analyzed in the primary screen, and the Z’ factors of these plates were between 0.5 and 0.8, showing that the results of our primary screen were reliable **(Figure 2B)**. In this study, we tested 3744 compounds at a final concentration of 10 μM to examine their capacity to inhibit the *Ld*PEX3-*Ld*PEX19 interaction. Hits from the primary screen were selected using a robust Z score. The robust Z score is insensitive to outliers because it is calculated with median and median absolute deviation (65, 66). Using robust Z score ≤ -3 as a threshold, which indicates significantly lower binding than the majority, 156 compounds were selected as primary hits **(Figure 2C)**. After the primary screen, we employed a counter screen assay to remove false positive hits. The signal drops observed in the primary screen might be caused by the effect of potential inhibitors on beads or energy transfer. Compounds might directly absorb the energy or fluorescence from beads or disrupt the interactions between protein tags and beads. Therefore, recombinant GST-His6 complex was used for the counter screen to exclude compounds that are false positive hits.

**Figure 2.**
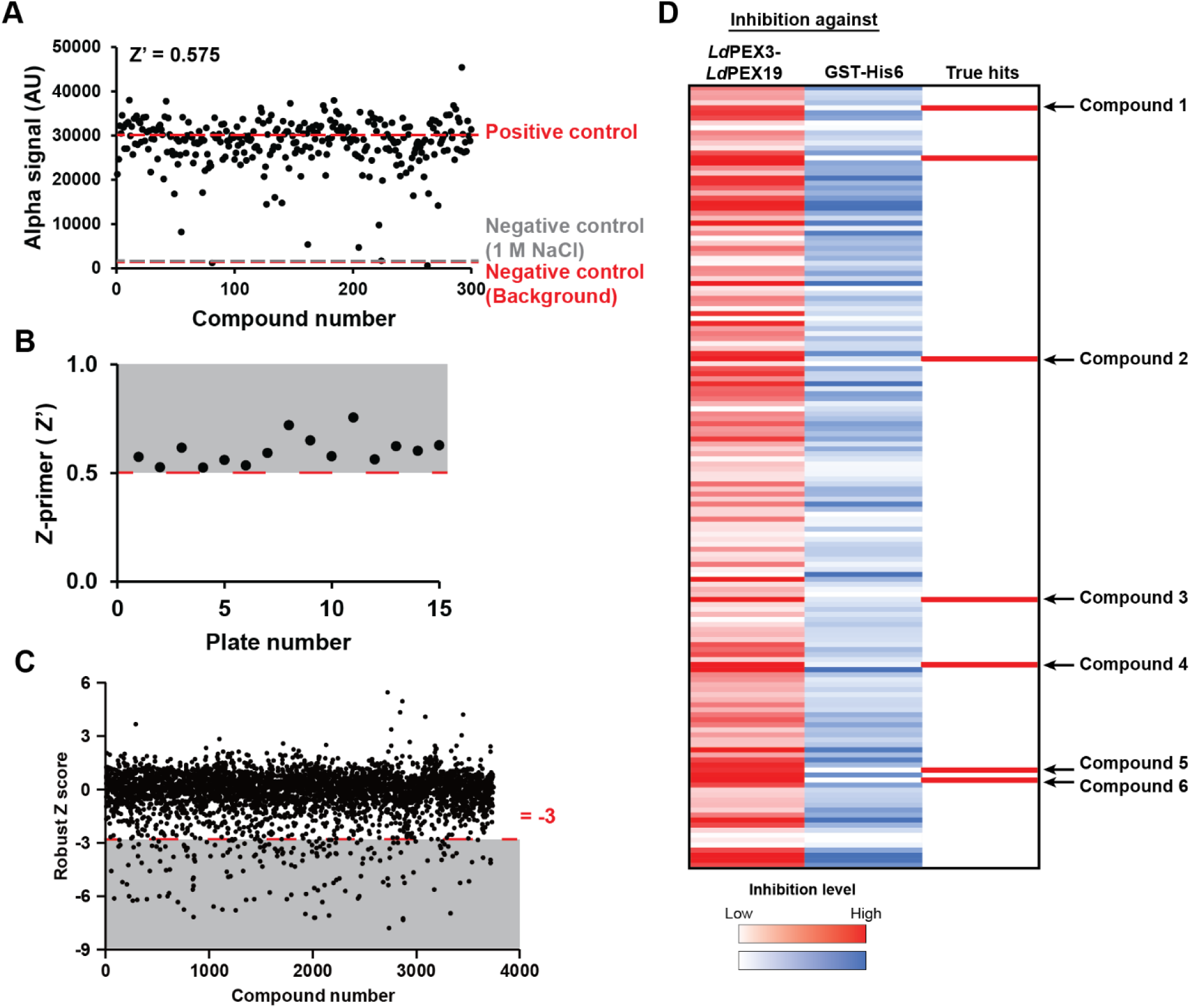
Screening of the repurposing drug library to identify inhibitors of *Leishmania* PEX3-PEX19 protein-protein interaction (PPI). **A)** The AlphaScreen based inhibitor screening was performed in 384-well plates, with each plate containing positive control (including both proteins and beads) and a negative control (beads only), which displayed normal protein interaction signal and background signal, respectively. To dissociate the protein complex as another negative control, 1M NaCl was used. The original Alpha signals from one plate are shown, with the average values of the controls indicated by dotted lines. The results demonstrate that only a few compounds affected the *Ld*PEX3-19 interaction. **B)** The Z-prime (Ź) factor was calculated from the positive and negative (NaCl added) signals to assess the assay quality of each plate. The figure shows that the Ź values for all plates were above 0.5, indicating that the screening results are reliable. **C)** A total of 3744 compounds from the drug repurposing library (MCE-HY-L035P-30 PartA, MedChemExpress) were tested for their ability to disrupt the *Ld*PEX3-PEX19 PPI at 10 μM. The compounds were separately added to the protein complex (8 nM) to evaluate their inhibitory effects. The robust Z-scores of all compounds are shown, with scores ≤ −3 (represented as dots in the grey region) shortlisted for counter-screening. **D)** Heatmaps in red and blue display the inhibitory effects of compounds on *Ld*PEX3-PEX19 or GST-His6 PPI, respectively. Darker red or blue indicates a stronger disruption of the interaction. The dark red squares, which overlap with dark red and light blue, represent six candidates that specifically inhibit the *Ld*PEX3-PEX19 PPI. One dark red square was excluded from further investigation as subsequent tests showed that it also inhibited the GST-His6 interaction (data not shown). AU: arbitrary units.

Compared to the *Ld*PEX3-*Ld*PEX19 complex, only half the concentration (4 nM) of the GST-His6 complex was used for the counter screen. This was based on the protein titration results **(Figure S2C)** because the signals of the GST-His6 complex were nearly double those of the *Ld*PEX3-*Ld*PEX19 complex. Therefore, we used 8 nM of *Ld*PEX3-*Ld*PEX19 but 4 nM of GST-His6 for the confirmation and counter screens, and primary hits were tested at 10 μM with protein complexes. Signals were normalized to the positive control (without compound treatment) and displayed as relative inhibition levels. In **Figure 2D**, the red heatmap displays the inhibition effects of compounds against the *Ld*PEX3-*Ld*PEX19 complex, and the intensity of the color indicates stronger inhibition of the protein-protein interaction (PPI). The blue heatmap depicts the apparent inhibition seen with GST-His6 protein, and the intensity of the blue color indicates the degree to which compounds affect the signal with the counter protein complex. From the counter screen, six compounds that specifically inhibited *Ld*PEX3-*Ld*PEX19 PPI (> 65 % inhibition) and did not inhibit signal with GST-His6 protein (< 10 % inhibition) were chosen as true hits for further characterization **(Figure 2D)**.

The shortlisted six compounds, denoted as Compounds 1, 2, 3, 4, 5, and 6, were estimated for their IC50 (half maximal inhibitory concentration) values using the dose-dependent assay against *Ld*PEX3-*Ld*PEX19 as well as GST-His6. In the previous screens, only one concentration (10 μM) was tested. Subsequently, a dose-dependent assay was performed with candidate compounds to assess their inhibitor activity. The IC50 values of Compounds 1 to 6 were 33.89 μM, 33.18 μM, 23.31 μM, 6.65 μM, 8.31 μM, and 7.91 μM, respectively **(Figure 3)**. No inhibition was detected with the GST-His6 complex (data not shown). For evaluation of specific inhibition, compounds were also tested with human proteins. GST-*Hs*PEX3Δ1-33 and *Hs*PEX19-His6 **(Figure S1F and S1G)** were performed in a cross-titration assay **(Figure S3A)**, and 10 nM of GST-*Hs*PEX3(Δ1-33) and 30 nM of *Hs*PEX19-His6 were used for the dose-dependent assay. The IC50 of Compound 1 in the *Hs*PEX3-*Hs*PEX19 complex was 15.85 μM, which is nearly half the concentration with *Leishmania* proteins **(Figure S3B)**. The IC50 values of Compounds 2 to 6 in human proteins were also estimated **(Table 2)**. Different from our expectations, compounds also inhibited the *Hs*PEX3-*Hs*PEX19 interaction, and some even showed stronger inhibition on human proteins. Therefore, compounds were further tested for cytotoxicity in human cells in parallel with anti-leishmanial activity assessment.

**Figure 3.**
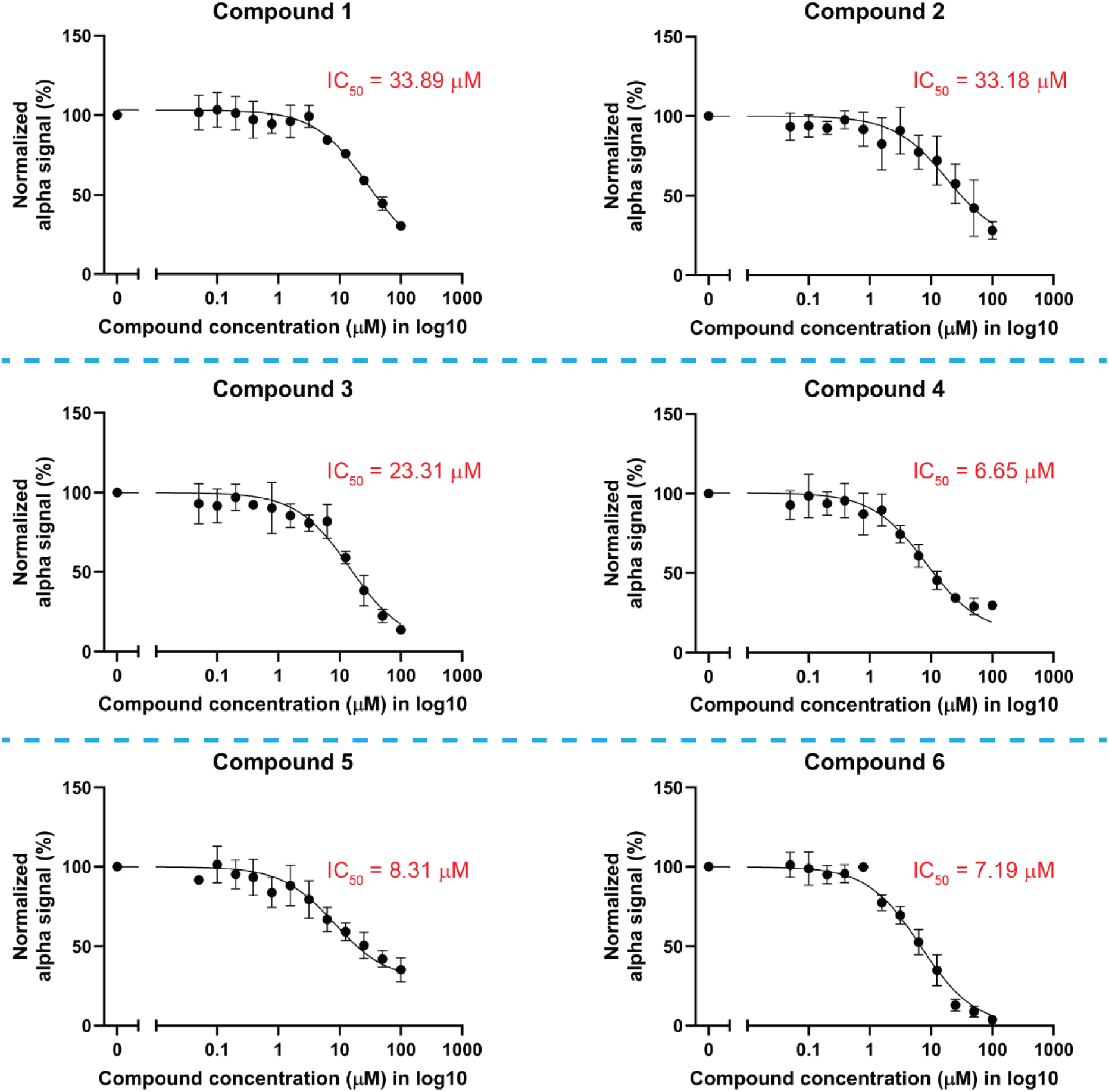
Dose-response curves for inhibition of the *in vitro* interaction of *Ld*PEX3-PEX19. Six shortlisted compounds were tested for their ability to inhibit the *Ld*PEX3-PEX19 interaction using an AlphaScreen based assay, with each compound tested at 12 concentrations (2-fold serial dilutions starting from 100 μM). The control signal (tagged proteins in the absence of any compounds) was set at 100% to normalize the results. The IC_50_ (half-maximal inhibitory concentration) values for Compounds 1 to 6 were determined to be 33.89 μM, 33.18 μM, 23.31 μM, 6.65 μM, 8.31 μM and 7.19 μM, respectively. The results of Compounds 1, 4 and 5 were obtained from three independent experiments, while those of Compounds 2, 3 and 6 were obtained from four independent experiments. Each data point represents the mean of replicates, and the error bars indicate the standard deviations. Data were plotted using GraphPad Prism 10, with curves fitted using the dose-response-inhibition mode (inhibitor vs. response, three parameters).

**Table 2.**

| Compound | *in vitro* activities against protein-protein interaction |  | Biological activities |  |  |  |
| --- | --- | --- | --- | --- | --- | --- |
|  | *Ld*PEX3-19<br>(IC<sub>50</sub>, μM) | *Hs*PEX3-19<br>(IC<sub>50</sub>, μM) | *L. tarentolae*<br>promastigotes<br>(EC<sub>50</sub>, μM) | HepG2<br>(TC<sub>50</sub>, μM) | Selective<br>index<br>(TC<sub>50</sub>/EC<sub>50</sub>) | *L. amazonensis* in<br>J774 macrophage<br>(EC<sub>50</sub>, μM) |
| 1 | 33.9 | 15.85 | 3.9 | 66.9 | 17 | 4.3 |
| 2 | 33.2 | > 100 | 22.1 | 66.4 | 3 | - |
| 3 | 23.3 | 44.4 | 16.4 | > 100 | > 6 | - |
| 4 | 6.7 | 2.23 | 14.6 | 91.5 | 6 | - |
| 5 | 8.3 | 26.5 | 53.1 | 19.7 | 0.4 | Not Tested |
| 6 | 7.2 | 15.5 | 23.4 | 99.6 | 4 | - |

### Anti-leishmanial activity and cytotoxicity in mammalian cells

To understand the killing effect of compounds, we tested the identified compounds for their toxicity to *L. tarentolae* promastigotes and human cells. To this purpose, we employed a resazurin-based assay to determine the EC50 (Half-maximal effective concentration, leading to a 50% reduction in the parasite survival) and TC50 (Half-maximal effective concentration, leading to a 50% reduction in the cell survival, i.e., cytotoxicity to human cells) in parasites and human cells, respectively. Compound 1 was the best anti-leishmanial candidate of all compounds, giving the EC50 value of 3.9 μM and a TC50 of 66.9 μM, with a selective index (ratio of TC50/EC50) of 17 **(Figure 4 and 5)**. Likewise, we also tested Compound 1 in other human cell lines, such as skin fibroblast GM05756, macrophage differentiated THP-1 and microglia HMC3. Considering HepG2 is a cancer cell line, it might show a higher tolerance to drug toxicity. Thus, GM05756 was tested for cytotoxicity as a normal cell line. Macrophage cell lines were chosen because *Leishmania* parasites reside in and multiply within macrophages. The results showed a low cytotoxicity of Compound 1 in the different human cells, with TC50 values above 70 μM **(Figure S3C - S3E)**. Compounds 2, 3, 4, and 6 were determined as effective anti-leishmanial compounds, with EC50 values of 22.1 μM, 16.4 μM, 14.6 μM and 23.4 μM, respectively **(Figure 4)**. Conversely, no apparent cytotoxicity was observed from these four compounds, with TC50 above 66 μM **(Figure 5)**. However, it was clear that Compound 5 was more toxic to human cells than parasites, given the TC50 of ∼20 μM but EC50 > 50 μM. Therefore, we only chose Compounds 1, 2, 3, 4, and 6 for further studies and mainly focused on Compound 1, considering its lower EC50 value and high selectivity.

**Figure 4.**
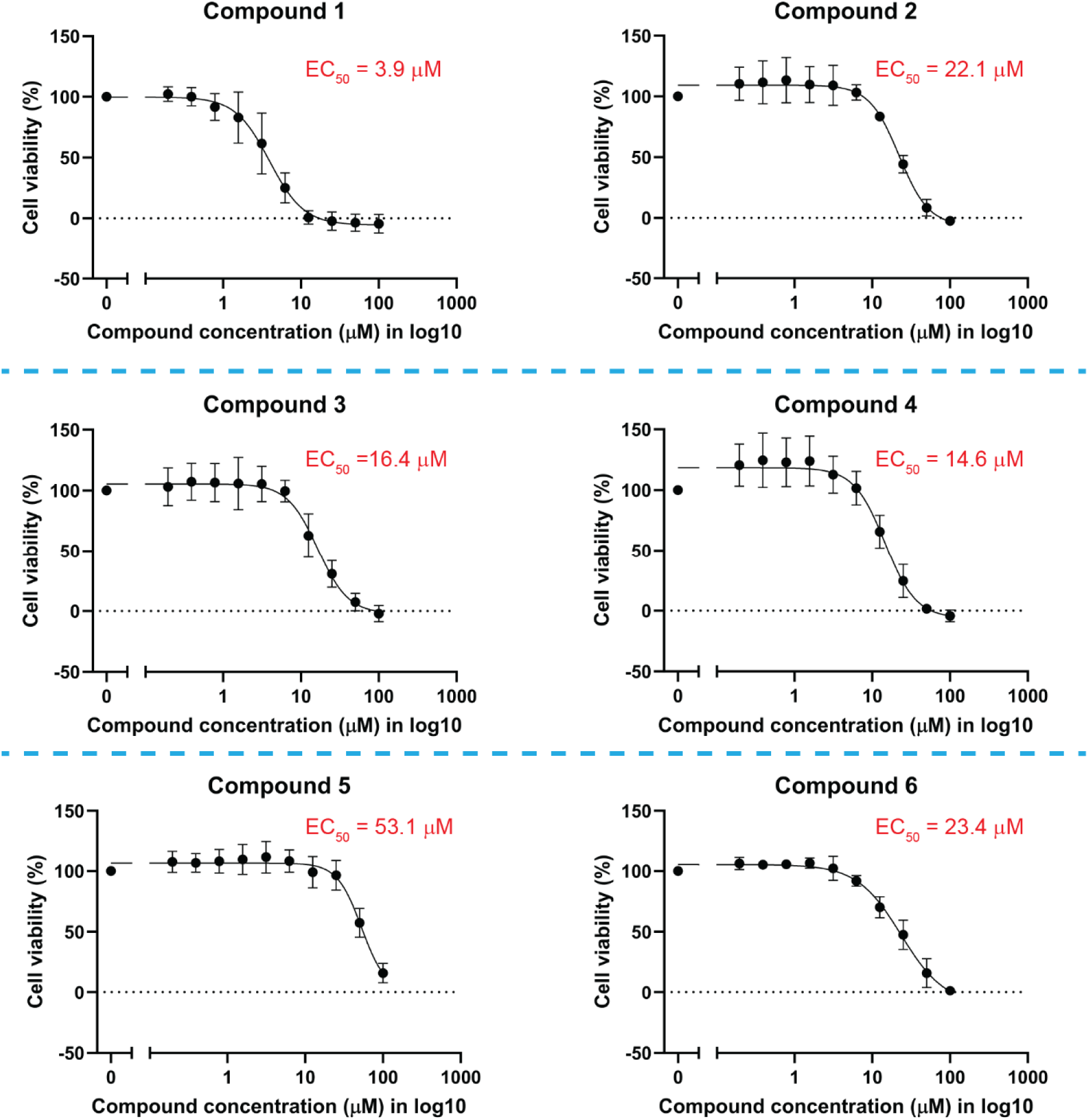
PEX3-PEX19 inhibitors show anti-leishmanial activity *in vitro*. Wild-type *Leishmania tarentolae* promastigotes (LtP) were treated with 2-fold serial dilution of inhibitors, starting at 100 μM. After 3 days of treatment, cell viability was assessed using a resazurin assay. The untreated control group was set as 100 % viability, and treated groups were normalized to this control. The EC_50_ (half maximal effective concentration) values of compounds were determined as follows: 3.9 μM (Compound 1), 22.1 μM (Compound 2), 16.4 μM (Compound 3), 14.6 μM (Compound 4), 53.1 μM (Compound 5), and 23.4 μM (Compound 6). The results were plotted and analyzed using a dose-response model (EC50 shift, X is concentration) in GraphPad Prism 10. Every data point represents the mean of three biological replicates, and the error bars indicate the standard deviations. Among the six compounds, Compound 1 exhibited the strongest anti-leishmanial effect, while Compound 5 was the least effective.

**Figure 5.**
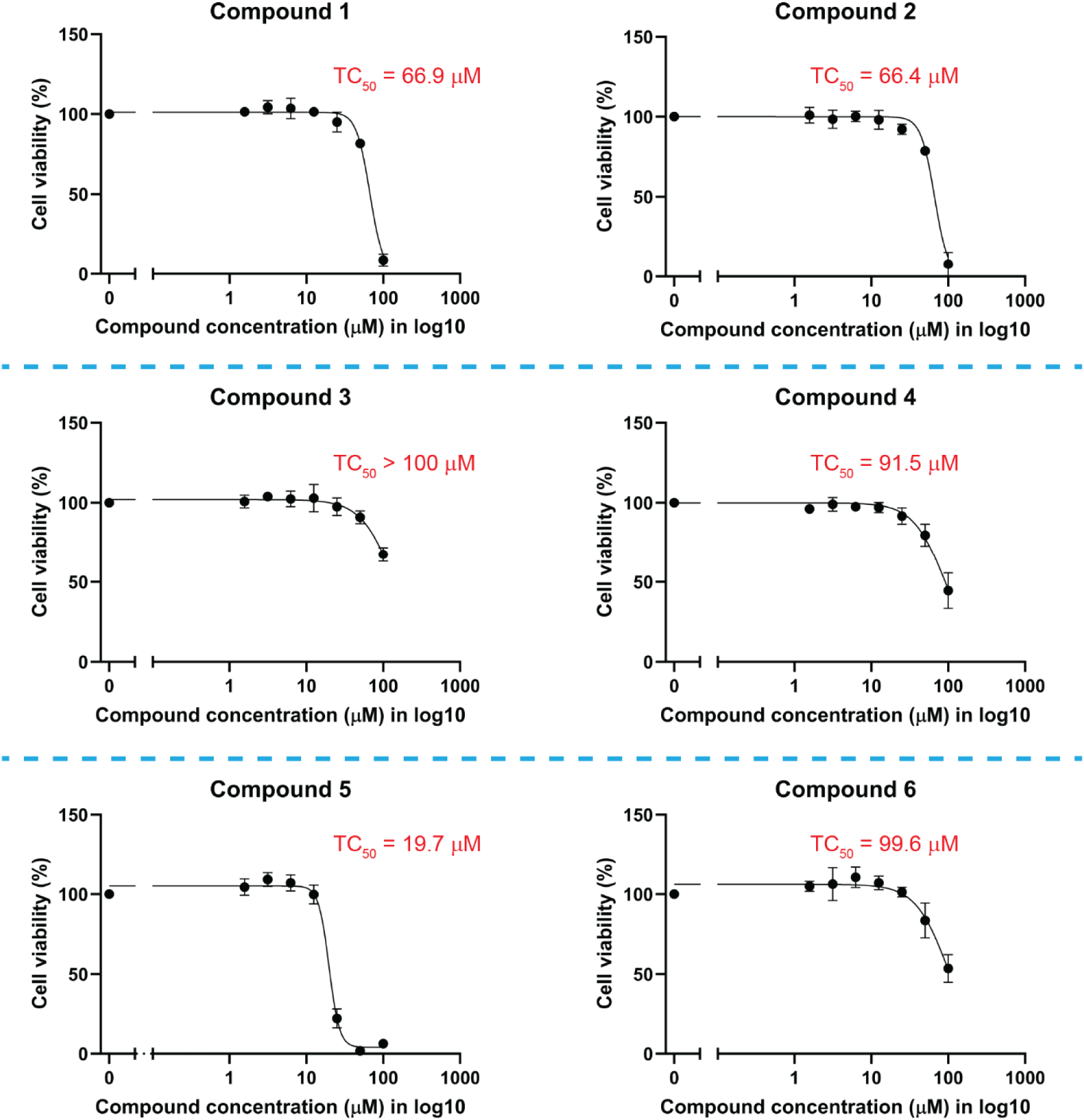
Cytotoxicity analysis of identified inhibitors against human cells. The human liver cancer cell line HepG2 was treated with six compounds, each serially diluted from 100 μM, to assess cytotoxicity. Cell viability was measured using a resazurin assay after being treated with compounds for three days. The viability of untreated cells was set at 100% and treated cells were normalized to this control. The TC_50_ (half-maximal toxic concentration) of Compound 5 was estimated as 19.7 μM, indicative for a significant drug toxicity to human cells. The other compounds exhibited low cytotoxicity against HepG2 cells, with TC_50_ values as follows: 66.9 μM (Compound 1), 66.4 μM (Compound 2), >100 μM (Compound 3), 91.5 μM (Compound 4), and 99.6 μM (Compound 6). Data were plotted with GraphPad Prism 10 and the curves were fitted using a dose-response model (EC50 shift, X is concentration). Each data point represents the mean of three biological replicates and the standard deviations are indicated with vertical error bars.

Furthermore, we assessed the activities of Compounds 1, 2, 3, 4, and 6 against *L. amazonensis* amastigotes inside infected murine macrophages using an *in vitro* cell-based assay (Performed by NYU Langone’s Anti-Infectives Screening Facility, USA). The EC50 of Compound 1 in this assay was 4.3 μM **(Figure 6A)**, and the TC50 was above 30 μM, resulting in a selective index of around 7. In comparison with the results depicted in **Figure 4**, we obtained a similar effective concentration in killing *Leishmania* parasites in different forms and species (*L. tarentolae* promastigotes and *L. amazonensis* amastigotes). Using other compounds, we did not detect any killing effect on the amastigotes **(Figure 6B)**. This may be a result of drug delivery problems. Since the amastigotes stay in membrane-bound vacuoles called *Leishmania* parasitophorous vacuoles (LPV) in the host macrophage (48, 73), they present an additional challenge in killing parasites because drugs need to permeate through multiple membranes, including the plasma membrane, phagosome, *Leishmania* cell membrane, and eventually reaching the glycosome. Consequently, one of our candidates, Compound 1, presents a potential inhibitor for anti-leishmanial treatment and should be further investigated in clinical research.

**Figure 6.**
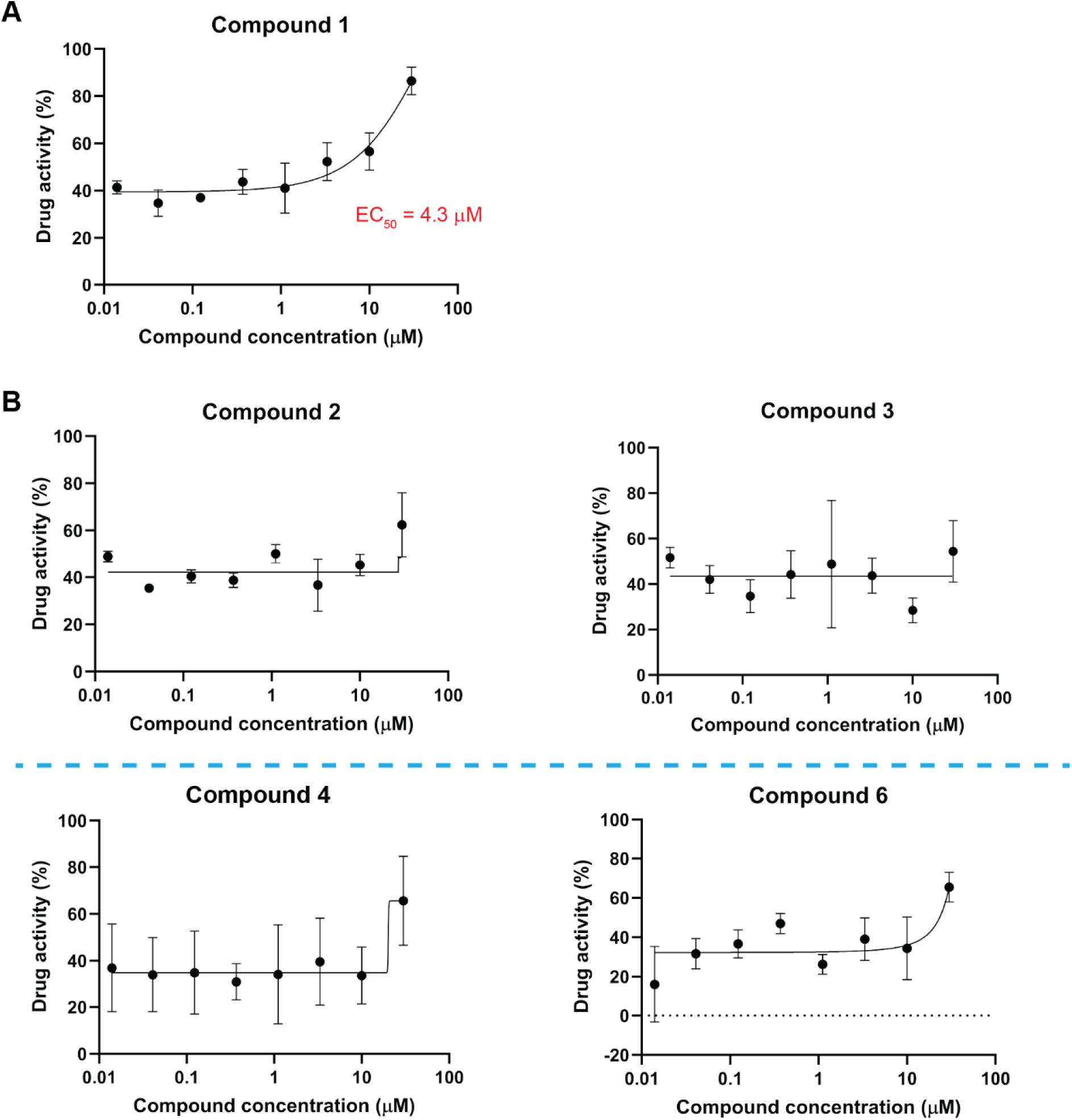
Anti-leishmanial activity analysis in a cell-based assay against *L. amazonensis* amastigotes. **A)** Compound 1 was evaluated for anti-leishmanial activity in a cell-based *in vitro* assay against human infective *Leishmania* species. Mouse macrophage (J774 cell line) cells were infected with *L. amazonensis* to establish the intracellular parasite stage called amastigotes. The infected cells were then treated with a serial dilution of Compound 1, starting from 30.3 μM. The resulting plot displays the drug activity, showing a lethal effect on parasites, with an EC_50_ value of 4.3 μM. **B)** Compounds 2, 3, 4, and 6 were also tested for anti-leishmanial activity against *L. amazonensis.* However, none of them exhibited considerable lethality in the assay, indicating they are not promising candidates for the development of anti-leishmanial treatments.

### Compound 1 specifically affects parasite glycosome biogenesis but not human peroxisomes

To understand how Compound 1 kills *leishmania* parasites, we applied a biochemical fractionation to investigate the glycosome matrix protein localization. *L. tarentolae* promastigotes were treated with Compound 1 or DMSO as an untreated control for 24 h. Then, cells were harvested and subjected to digitonin fractionation for the glycosomal protein mislocalization study. Six concentrations of digitonin varying from 0.02 to 1 mass ratio (digitonin/protein, w/w) were incubated with cells separately. With a low concentration of digitonin (0.025 mg of digitonin/mg of total protein), glycerol kinase (GK) was released into the supernatant fractions in the treated cells but not in the untreated cells **(Figure 7A)**. GAPDH and aldolase were also released at a lower concentration of digitonin (0.2 mg of digitonin/mg of total protein) in the treated cells, indicating that Compound 1 treatment causes the matrix protein to mislocalize to the cytosol. Cytosol protein enolase was released at lower concentrations of digitonin, showing that cell membranes were permeabilized. However, we found that the enolase of treated cells was released in higher quantities than untreated, while we expected the enolase to be released at the same level. We guess this was because parasites treated with Compound 1 were very fragile, and cell membranes were totally broken at low concentrations of digitonin, leading to high quantities of enolase release. Tubulin was released when 0.1 mg digitonin was applied to untreated parasites. The releasing pattern of tubulin is similar to enolase since the sub-cellular location of tubulin is also cytosol. mtHSP70, as a mitochondria matrix protein (74), was used to evaluate the effect on mitochondria. Nearly no mtHSP70 was released from the parasites, representing that Compound 1 might not significantly affect the mitochondria. In parallel, cells treated with Compound 1 or DMSO were fixed, permeabilized, and immunolabeled using antibodies against glycosomal marker enzyme aldolase. Immunofluorescence images revealed red punctata, which represent the organellar aldolase localization in the cells **(Figure 7B)**. Compared to the DMSO control, there were larger red puncta in the compound-treated cells. Usually, there exist many smaller glycosomes in cells that can be detected with aldolase labeling. However, different patterns of glycosome were observed when Compound 1 was added to cells. The different patterns indicate that glycosomes might be enlarged or cluster due to Compound 1 treatment, which could be related to an inhibition of glycosome biogenesis. These results demonstrate that Compound 1 indeed has an effect on glycosome but might also affect other organelles. The observed killing effect of Compound 1 on parasites might be caused by disturbing glycosome biogenesis and other metabolic pathways.

**Figure 7.**
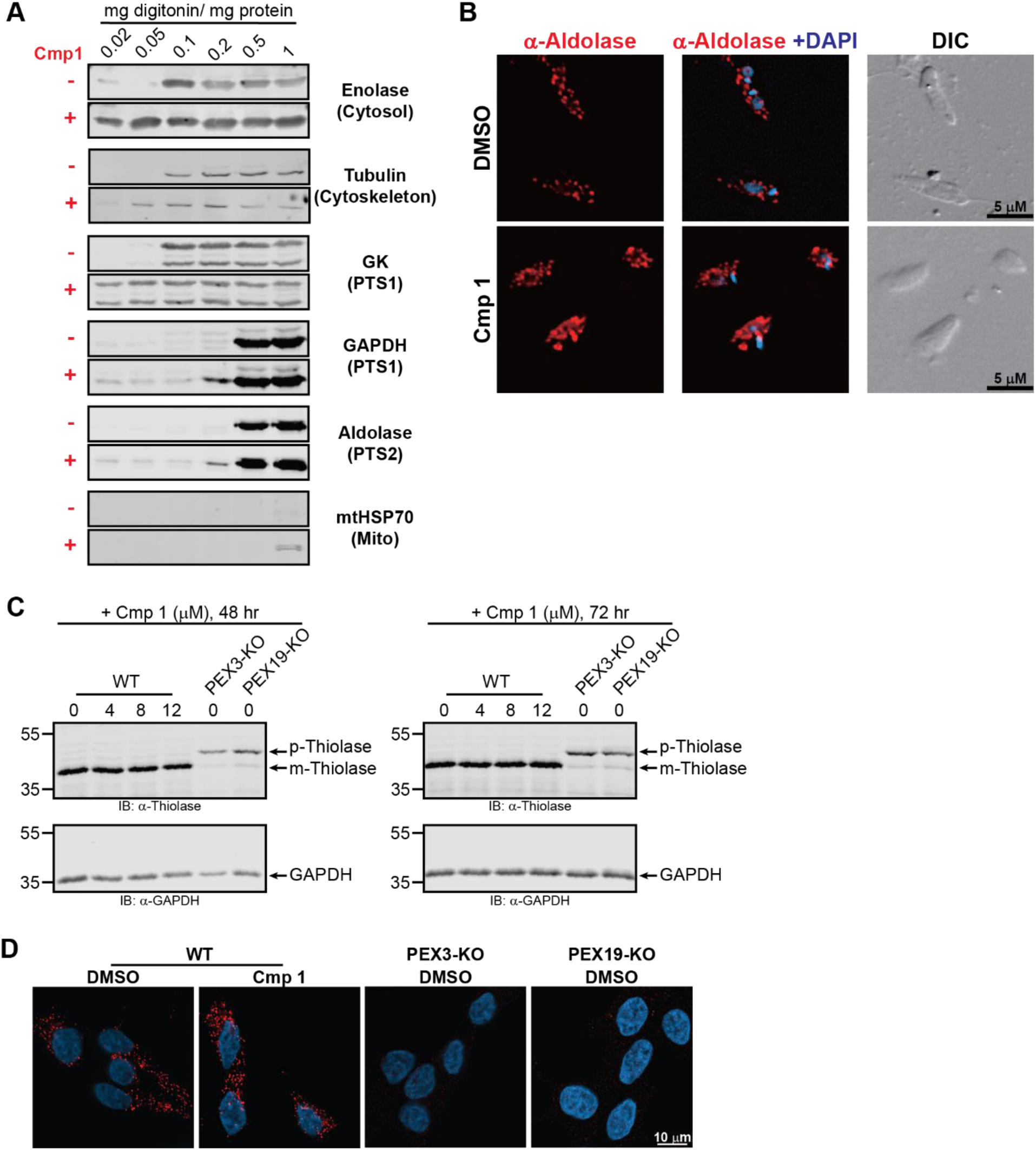
Analysis of the effects of Compound 1 on glycosomes/ peroxisomes (Target validation). **A)** Wild-type *L. tarentolae* promastigotes (LtP) were treated with either DMSO(-) or 12 μM of Compound 1 (+) for 24 h. Digitonin fractionation was performed to investigate the mislocalization of glycosome matrix proteins. Enolase was used as cytosolic marker. The PTS1 proteins glycerol kinase (GK) and glyceraldehyde 3-phosphate dehydrogenase (GAPDH) served as glycosome matrix markers for PTS1 import, while aldolase served as a marker for glycosomal PTS2 import. The mitochondrial heat shock protein (mtHSP70) was used as a mitochondrial matrix marker to confirm that mitochondria were unaffected. Compared to the untreated group, GK, GAPDH, and aldolase were released from Compound 1-treated cells at lower levels of digitonin. The result indicated a partial mislocalization of glycosomal matrix proteins to cytosol upon Compound 1 treatment. **B)** LtP cells treated with DMSO or Compound 1 were fixed after 24 h of incubation and stained with anti-*Tb*Aldolase primary antibody. The red punctuate staining indicates the glycosomal localization of aldolase. Compared to the DMSO control group, cells treated with Compound 1 showed bigger red puncta, which might be related to enlarged or cluster glycosomes, suggesting a mild effect of Compound 1 on glycosomes. **C)** Human T-REx-293 cells were incubated with three concentrations of Compound 1 for 48 or 72 h and analyzed via immunoblotting using 3-ketoacyl-CoA thiolase (Thiolase) primary antibody. Precursor thiolase (p-Thiolase) is proteolytically processed upon peroxisomal import into its mature form (m-Thiolase), which runs at 44 kDa and serves as an indicator for peroxisomal import. Compound 1-treated wild-type (WT) cells displayed an m-Thiolase pattern similar to DMSO-treated cells, indicating that the peroxisomal protein import is normal. PEX3- and PEX19-knockout (KO) cells, in which p-Thiolase is abundant due to the absence of peroxisomes, were used as controls. **D)** Human T-REx293 cells, PEX3-KO, and PEX19-KO cells were grown on coverslips and treated with DMSO or 12 μM of Compound 1 for 72 h. Peroxisomes were labeled using anti-PMP70 antibody, whereas nuclei were stained with DAPI. Compound 1-treated cells showed a peroxisome pattern like that of the control (DMSO-treated) cells, indicating normal peroxisome morphology. No peroxisomes were observed in PEX3-KO and PEX19-KO cells.

Because Compound 1 also inhibited the *Hs*PEX3-*Hs*PEX19 interaction in the previous test, it was tested with human T-Rex293 cells to evaluate its effects on human peroxisomes. Cells were treated with Compound 1 for 48 or 72 h, and then the whole cell lysates were subjected to immunoblot analysis. Thiolase, a PTS2 protein, was used in this study as an indicator for peroxisomal protein import (75, 76). Pre-mature thiolase is synthesized in the cytosol and imported into peroxisomes where proteolytic cleavage of the PTS2 takes place resulting in mature thiolase (77, 78). The mature thiolase can be distinguished from the pre-mature thiolase based on the molecular weight difference due to the PTS2 cleavage. The migration shift can be assessed in the immunoblot. Furthermore, the dominant pre-mature thiolase is detected when peroxisomes are defective or absent. Here, PEX3 and PEX19 CRISPR knockout (KO) cells were applied as negative controls, as peroxisomes are either defective or absent. In **Figure 7C**, a dominant band of mature thiolase was detected in the compound-treated and untreated wild-type cells. In comparison, pre-mature thiolase was the majority in PEX3-KO and PEX19-KO cells. Using peroxisome membrane protein PMP70 as a peroxisomal marker, fluorescence microscopy revealed that clearly normal peroxisomes were found in compound-treated cells but not in PEX3-KO and PEX19-KO cells **(Figure 7D)**. With these results, we validated that Compound 1 does not affect the PTS2 import machinery or peroxisome morphology in human cells. In summary, we confirmed that Compound 1 affects glycosome biogenesis in *Leishmania* cells but does not affect human peroxisomes.

## Discussion

More than one million new patients suffer from leishmaniasis every year, and over one billion people are at risk of infection (79). However, effective treatments remain limited because of side effects, such as drug toxicity and unfordable costs (80, 81). Additionally, a growing number of resistant strains of *Leishmania* species have appeared, hindering progress in leishmaniasis treatment. This underscores the urgent need to develop new drugs for leishmaniasis. Previous studies have shown that glycosomes, which compartmentalize glycolytic and other metabolic enzymes in the biological pathway, are crucial for parasite survival (82, 83). It was found that targeting hexokinase to glycosomes is essential to prevent cytotoxicity from uncontrolled glucose phosphorylation in *L. donovani* (84). In addition, the purine salvage pathway is critical in *Leishmania* due to the absence of a *de novo* purine biosynthesis pathway. The enzymes involved in purine and pyrimidine metabolism are also compartmentalized in glycosomes (85). Therefore, disrupting glycosome biogenesis represents a potential strategy for leishmaniasis treatment.

Previous research revealed that glycosomal enzymes, as well as glycosomal biogenesis, are promising drug targets (55, 86) and showed that small molecule inhibitors could block the import of glycosomal matrix proteins (87). The inhibitors of PEX5-PEX14 indicated therapeutic effect in animal models infected by *T. brucei* (60). Furthermore, another two studies demonstrated the potential to kill *T. brucei* by disrupting glycosomal membrane protein biogenesis (61, 62). Since blocking PEX3-PEX19 interaction can directly affect glycosomal membrane protein import as well as glycosomal matrix protein import, inhibitors of these processes likewise are promising tools for novel therapies. Moreover, the sequence similarity of PEX3 between humans and *Trypanosoma* or *Leishmania* PEX3 is very low, which indicates a reduced chance of off-target effect. Recently, inhibitors of *L. donovani* PEX5-PTS1 were identified using fluorescence polarization-based screening (88). These studies reveal that inhibitors for disrupting glycosome biogenesis could effectively treat trypanosomatid diseases. In the past, most researchers focused on blocking glycosome matrix protein import. However, in recent years, interest and efforts have grown in the membrane protein import machinery, and studies have increasingly targeted the function of glycosomes, thus addressing the problem of trypanosomiasis at their root. Therefore, we transferred these strategies to the *Leishmania* system because *Leishmania* species share a similar PEX3 sequence to *T. brucei*, implying that hindering the interaction between the membrane protein PEX3 and the corresponding cytosolic PEX19 protein could be an effective treatment in leishmaniasis.

In this study, we established the AlphaScreen based assay using recombinant *Ld*PEX3 and *Ld*PEX19 proteins. In the saturation binding assay, we confirmed the interaction between these two proteins, and the capability of disrupting the interaction was shown in the competition binding assay. A drug repurposing library was applied in this study for drug screening. In this decade, a growing number of repurposed drugs have been tested in different studies due to their benefits. These compounds might readily be available to patients with unrelated diseases because they either already exist on the market or are in clinical trials and have, therefore, passed toxicity tests. Here, we found six candidate compounds from the library screening and confirmed their inhibition in dose-dependent assays with IC50 values ranging from 7 – 34 μM. Via dose-dependent assays, Compounds 4 to 6 showed better inhibition compared to Compounds 1 to 3. Concerning the killing effect on *L. tarentolae* promastigotes, the EC50 of Compounds 5 was above 50 μM, which we did not consider to be efficacious. Compounds 2, 3, 4, and 6 exhibited EC50 between 14 -23 μM. No apparent cytotoxicity for Compounds 2, 3, 4, and 6 was seen. However, these four compounds also were not effectively killing *L. amazonensis* amastigotes.

The IC50 value for inhibition of the *Ld*PEX3-19 interaction by Compound 1 was higher than for the inhibition of the *Hs*PEX3-19 interaction. Nevertheless, Compound 1 was identified as the best candidate from this library, because of the EC50 of ∼4 μM against *L. tarentolae* promastigotes as well as *L. amazonensis* amastigotes, while displaying minimal cytotoxicity to human cells. The selective index of Compound 1 for *L. tarentolae* promastigotes is above 15 and ∼7 was for *L. amazonensis* amastigotes. DNDi guidelines indicate that the selectivity index should be a minimum of 10 for early hits on screens (89), indicating the suitability of the hit for further optimizations. Even though the inhibitory effect was stronger in human proteins than in *Leishmania* proteins, the killing effect was the opposite in human cells and parasites. The results might be due to the unique characteristics of glycosomes, as they are essential for parasite survival, while peroxisomes are not for human cells. Moreover, when treated with Compound 1, glycosomal proteins were mislocalized in promastigotes, but human peroxisomes were unaffected. The mitochondria matrix protein (mtHSP70) was slightly released in Compound 1 treated parasites with a high concentration of digitonin, and this might indicate that mitochondria also be affected by Compound 1. This data suggests that Compound 1 mediates its killing efficiency by inhibiting protein import into glycosomes; however, we cannot exclude that the drug also affects other essential pathways.

In conclusion, we found that the described candidate compounds can disrupt the interaction between *Ld*PEX3 and *Ld*PEX19 using the AlphaScreen assay. Compound 1 was identified as a potential inhibitor against various *Leishmania* species. Further investigation of Compound 1’s efficacy in animal models is necessary and urgent. Our findings indicate that blocking the PEX3-PEX19 interaction is a suitable approach against *Leishmania* species and provides a promising alternative treatment for leishmaniasis.

## Author contributions

SC, RE, and VK conceived and planned the experiments. SC performed all experiments, with RE and VK supervision. SC wrote the manuscript with support from RE and VK. All authors read and gave feedback on the manuscript.

## Acknowledgements

This work was supported by the Deutsche Forschungsgemeinschaft (ER 178/17-1) to RE, and InnovationsFoRUM grants of the Ruhr-University Bochum IF-009N-22, IF-018N-22) to RE. We thank Prof. Dr. Paul Michels for kindly providing various *Trypanosoma* antibodies. We thank Dr. Chethan Kumar Krishna for his suggestions about this study and for reading the manuscript. We thank Samantha Jeng for reading the manuscript. We thank Bettina Tippler for assisting in drug dilution and Nadine Schmidt for assisting in microscopy.

---

**Supplementary figure 1.**
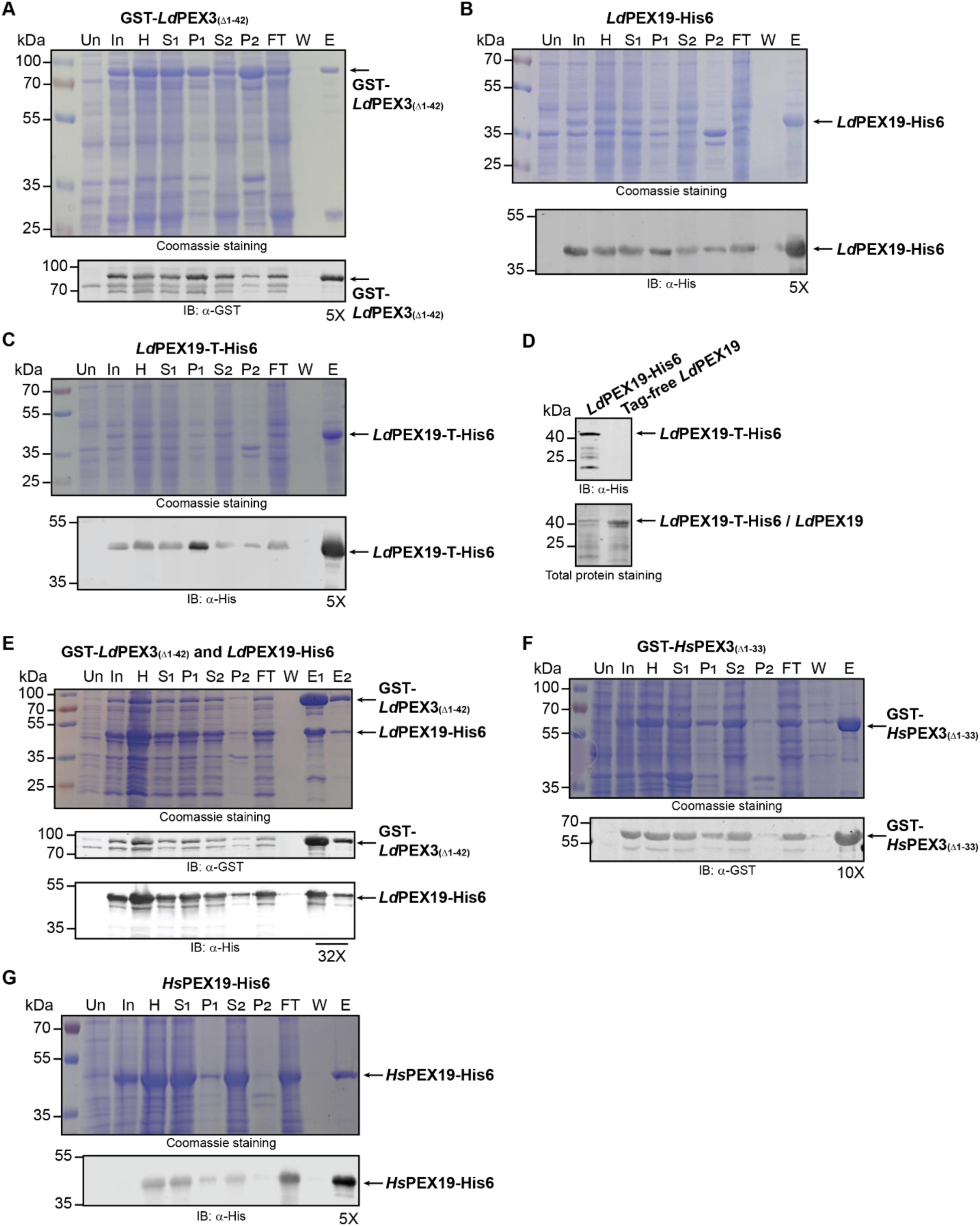
Protein purification profiles. The protein purification profiles shown in A-C and E-F were analyzed by SDS-PAGE followed by Coomassie staining (upper panels) and immunoblotting (lower panels) with antibodies against GST or the His6 tag as indicated. **A)** The recombinant GST-*Ld*PEX3_Δ1-42_ protein (80 kDa) was expressed in *E. coli* and purified by affinity chromatography on glutathione agarose beads. **B)** The C-terminally His6-tagged full-length *Ld*PEX19 protein (35 kDa) was purified by affinity chromatography on Ni-NTA agarose beads and subsequently used for the AlphaScreen based assay. **C)** *Ld*PEX19 fused with a thrombin cleavage site and a C-terminal His6 tag (*Ld*PEX19-T-His) was purified by affinity chromatography on Ni-NTA agarose beads. A thrombin cleavage site (T) was inserted between full-length *Ld*PEX19 and the His6 tag, resulting in 36-kDa fusion proteins. **D)** The tag-free *Ld*PEX19 proteins were generated from recombinant *Ld*PEX19-T-His6 protein by reacting with thrombin enzyme. Ni-NTA agarose beads were used to remove cleaved His6 tag and any remaining uncleaved portion of *Ld*PEX19-T-His6. Both *Ld*PEX19-T-His6 and tag-free proteins were analyzed using immunoblotting with anti-His antibody and total protein staining (Revert Total Protein Stain 700, LI-COR). **E)** GST-*Ld*PEX3_(Δ1-42)_ and *Ld*PEX19-His6 were co-expressed and co-purified by affinity chromatography on glutathione agarose beads. The elution fraction contained both GST-*Ld*PEX3_(Δ1-42)_ and *Ld*PEX19-His6 proteins, which were subsequently analyzed by immunoblotting with anti-GST and anti-His antibodies. **F)** The recombinant GST-*Hs*PEX3_(Δ1-33)_ protein, lacking the predicted transmembrane domain in the N-terminus, was expressed in *E. coli* and purified by affinity chromatography on glutathione agarose beads. The GST-*Hs*PEX3, predicted to be 63 kDa, is indicated with an arrow. **G)** *Hs*PEX19-His6 (35 kDa) was purified by affinity chromatography on Ni-NTA agarose beads for the cross-titration and dose-response curve assays with GST-*Hs*PEX3_(Δ1-33)_ proteins. Un: uninduced sample, In: induced sample, H: homogeneous fraction, S_1_: first supernatant fraction, P_1_: first pellet fraction, S_2_: second supernatant fraction, P_2_: second pellet fraction, FT: flow-through, W: wash, E: elution fraction. E_1_: first elution fraction, E_2_: second elution fraction. Numbers listed below elution fractions such as 5X, 32X and 10X indicate the concentration factors.

**Supplementary figure 2.**
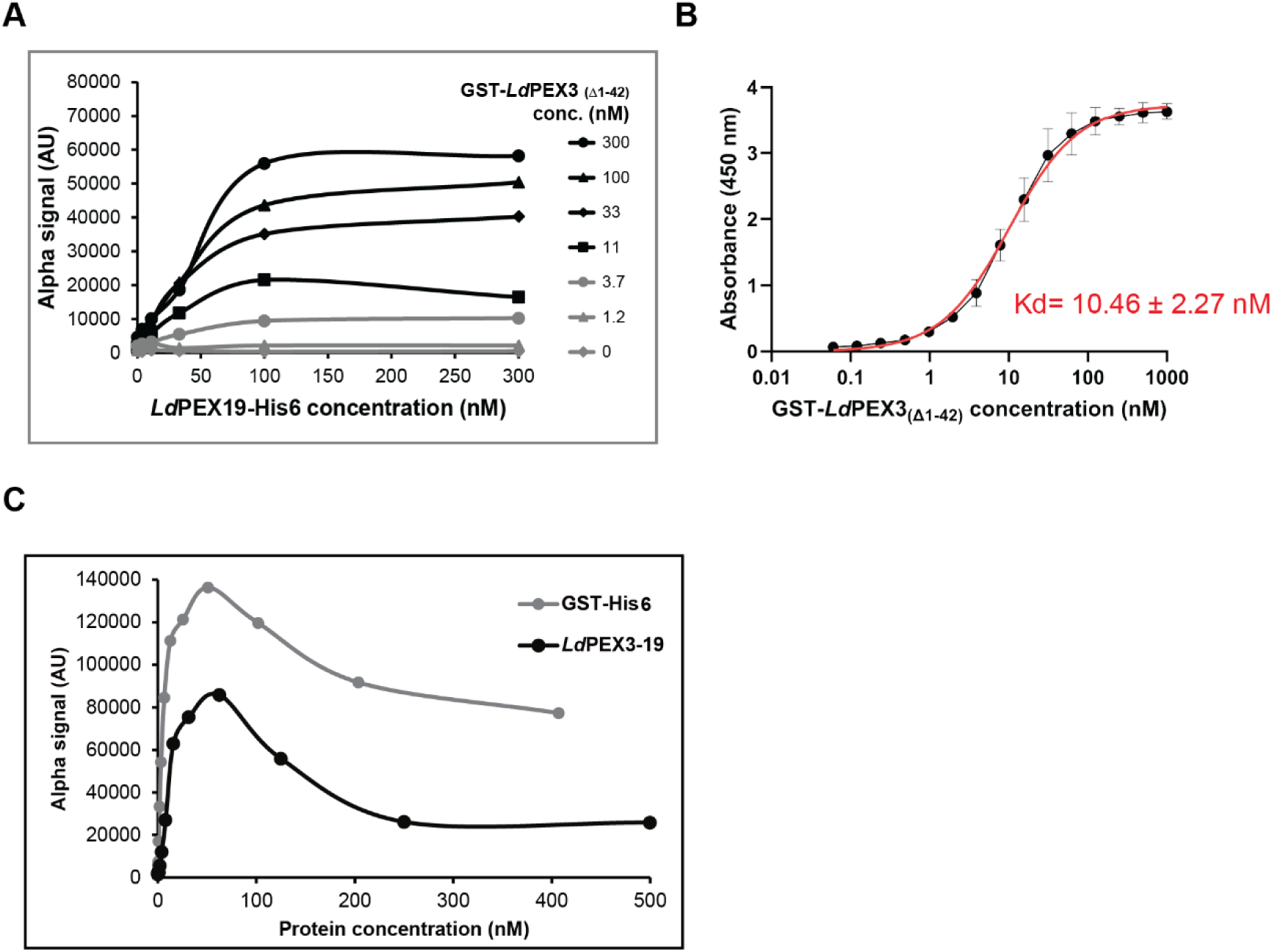
*Ld*PEX3 interacts with *Ld*PEX19 in AlphaScreen based and ELISA assays. **A)** Cross titration of GST-*Ld*PEX3(Δ1-42) protein and *Ld*PEX19-His6 protein was performed in the AlphaScreen based assays up to a concentration of 300 nM. Recombinant GST-*Ld*PEX3(Δ1-42) and *Ld*PEX19-His6 proteins were incubated at room temperature. Subsequently, acceptor beads and donor beads were added separately to interact with the proteins. Results are expressed in arbitrary units (AU). **B)** *Ld*PEX19-His6 proteins were first coated to the 96-well plate, and unbound proteins were washed away. GST-*Ld*PEX3(Δ1-42) proteins were then added to the pre-coated wells to interact with the *Ld*PEX19-His6 proteins. Anti-GST antibody (1:1000) was used to detect GST-*Ld*PEX3(Δ1-42) proteins and anti-mouse horseradish peroxidase (1:1000) was used to amplify the signal. Chromogenic substrate 3,3’,5,5’-tetramethylbenzidine (TMB) was added to develop color, and 0.5 M H_2_SO_4_ was used to stop the color development. The dissociation constant (Kd) was estimated as 10.46 ± 2.27 nM. Data were collected from three replicates and analyzed using GraphPad Prism 10. The curve was fitted with the one site-specific binding model and each data point represents the mean. Error bars indicate the standard deviations. **C)** The co-expressed and co-purified GST-*Ld*PEX3(Δ1-42) -*Ld*PEX19-His6 protein complex and GST-His6 protein complex were tested in the AlphaScreen based assay. Both protein complexes were serially diluted and incubated with acceptor and donor beads for signal detection. The maximum signals for both protein complexes were observed at a concentration of approximately 60 nM.

**Supplementary figure 3.**
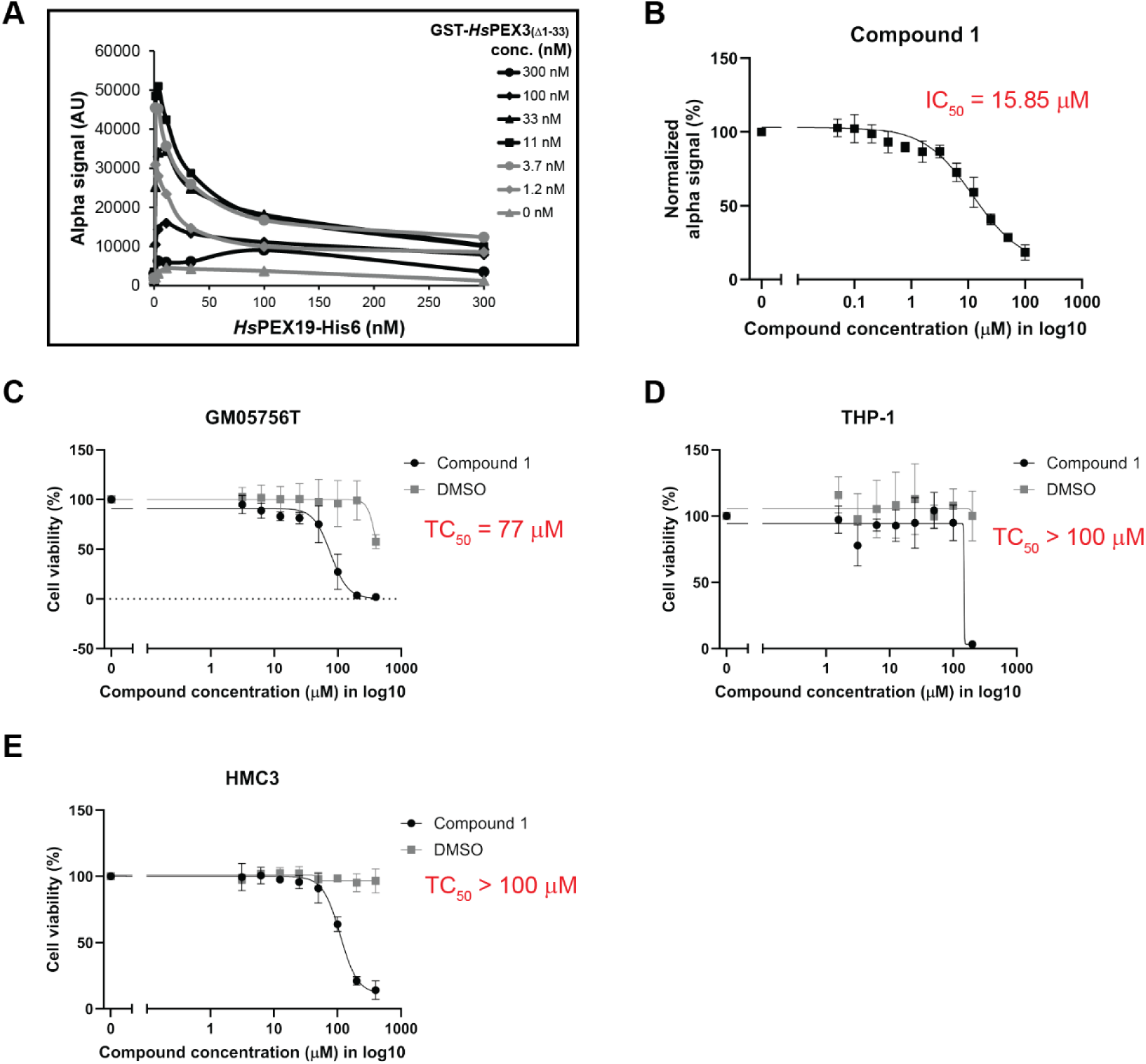
Characterization of Compound 1 effects on *Hs*PEX3-PEX19 interaction and on the survival of human cell lines. **A)** Cross titration of GST-*Hs*PEX3_(Δ1-33)_ and *Hs*PEX19-His6 proteins was performed using 2-fold serial dilutions starting from 300 nM in the AlphaScreen based assays. The maximum signal (∼50000 AU) for the interaction between two proteins was detected when GST-*Hs*PEX3_(Δ1-33)_ was at 11 nM and *Hs*PEX19-His6 was at 3.7 nM. AU: arbitrary units. **B)** Compound 1 was tested in 2-fold serial dilutions starting from 100 μM against human PEX3-PEX19 in a dose-response curve analysis. The IC_50_ (half-maximal inhibitory concentration) of Compound 1 against the human proteins was determined to be 15.66 μM. Data were collected from triplicates and each data point represents the mean, with error bars indicating standard deviations. **C, D, E)** Compound 1 was tested for cytotoxicity in different human cell lines as indicated. The TC_50_ values were estimated to be 77 μM for GM05756T (skin fibroblast), and greater than 100 μM for both THP-1 (monocytes) and HMC3 (microglia). Combined with Figure 5, these results indicate no significant cytotoxicity for human cells of Compound 1 was observed. Cytotoxicity assays were independently repeated at least three times. Data were analyzed using curve fitting with a dose-response model (EC50 shift, X is concentration) in GraphPad Prism 10.

